# The naked cell: emerging properties of a surfome-streamlined *Pseudomonas putida* strain

**DOI:** 10.1101/2020.05.17.100628

**Authors:** Esteban Martínez-García, Sofía Fraile, David Rodríguez Espeso, Davide Vecchietti, Giovanni Bertoni, Víctor de Lorenzo

## Abstract

Environmental bacteria are most often endowed with native surface-attachment programs that frequently conflict with efforts to engineer biofilms and synthetic communities with given tridimensional architectures. In this work we report the editing of the genome of *Pseudomonas putida* KT2440 for stripping the cells of most outer-facing structures of the bacterial envelope that mediate motion, binding to surfaces and biofilm formation. To this end, 23 segments of the *P. putida* chromosome encoding a suite of such functions were deleted, resulting in the surface-naked strain EM371, the physicochemical properties of which changed dramatically in respect to the wild type counterpart. As a consequence, surface-edited *P. putida* cells were unable to form biofilms on solid supports and—because of the swimming deficiency and other physicochemical alterations—showed a much faster sedimentation in liquid media. Surface-naked bacteria were then used as carriers of interacting partners (e.g. Jun-Fos domains) ectopically expressed by means of an autotransporter display system on the now easily accessible cell envelope. Abstraction of individual bacteria as adhesin-coated spherocylinders enabled rigorous quantitative description of the multi-cell interplay brought about by thereby engineered physical interactions. The model was then applied to parameterize the data extracted from automated analysis of confocal microscopy images of the experimentally assembled bacterial flocks for analyzing their structure and distribution. The resulting data not only corroborated the value of *P. putida* EM371 over the parental strain as a platform for display artificial adhesins but also provided a strategy for rational engineering of distributed biocatalysis.

---

Although the fundamental and biotechnological value of designer microbial communities is beyond any doubt ^1-3^, the endeavor is clearly limited by the fact that naturally-occurring bacteria have their own surface-attachment agenda ^4-6^. This makes native adhesion properties and endogenous biofilm formation programs to interfere with efforts to engineer multi-strain interactions and/or programmable biofilms. The logical way forward is suppression of the innate mechanisms of surface adhesion and replacement by artificial counterparts. Since many types of bacteria control a switch between sessile and surface-attached lifestyles by regulating the intracellular levels of cyclic-di-GMP ^7-9^ it comes as little surprise that most efforts thus far to manage adhesion have attempted to override the endogenous cyclic-di-GMP regulon e.g. by heterologous expression of a specific cyclase ^10^. An alternative approach involves the display of heterologous adhesins on the bacterial surface—typically protein interacting partners thereof ^11^ or nanobodies/matching antigens ^12-14^— so that their conditional expression brings about more or less structured multi-cell assemblies. While this last strategy looks more promising for engineering bacterial communities with given tridimensional (3D) structures, it relies on mutual accessibility of the interacting partners that protrude from the outer cell membrane. This is often made difficult by the presence of a suite of bulky envelope components that impede direct contact between bacterial surfaces. The envelope of Gram-negative bacteria is characteristically composed of an inner membrane, a peptidoglycan layer and an asymmetrical outer membrane (OM) that is directly facing the environment. The OM is composed of polyanionic lipopolysaccharides and phospholipids in outer and inner layers which in turn hold lipoproteins, porins, gated-channels and exports systems ^15-16^. Yet, the OM is also the scaffold of a suite of non-lipidic structures on its outward-facing layer that allow interactions with the environment in different ways e.g. swimming to explore different niches (flagella), inter-cell adhesion or attachment to solid surfaces (pili, fimbria, surface proteins and exopolysaccharides; ^6, 17^). While these structures are important for enduring different environmental challenges, they may be otherwise dispensable for biotechnological applications ^18-19^ including the above-mentioned design of structured bacterial consortia ^20^.

In order to expand the utilities of *Pseudomonas putida* as a prime synthetic biology chassis for industrial and environmental uses, we set out to edit the genome of the reference strain KT2440 for making it an optimal biological frame for surface display of artificial adhesins. *P. putida* KT2440 is a non-pathogenic, HV1 certified ^21^, soil inhabitant and root colonizer endowed with a robust metabolism. Its diverse biochemical capacities enable this strain to use a broad number of substrates as carbon sources and host strong redox reactions ^22-24^. Moreover, it is highly resistant to organic solvents ^25^ and it is amenable to a large number of genetic manipulations ^26-28^. Taken together, these properties make *P. putida* KT2440 a platform of choice for bioproduction of value-added compounds ^29-31^. Alas, as is the case with other *Pseudomonas, P. putida* KT2440 has also a complex cell envelope with a number of structures that obstruct the access to proteins closely attached to the cell surface or artificially designed adhesins thereof.

In this work we have scanned the genome of *P. putida* KT2440 to identify non-essential genes encoding surface-protruding structures innately exposed on the OM. Most of them were serially deleted to generate cells with a less complex and more accessible outer cell surface. This operation resulted in an envelope-streamlined strain that was morphologically and phenotypically characterized and also tested as carrier of artificial adhesins displayed on the cell envelope by means of genetically encoded autotransporter domains ^11, 32^. The data below not only demonstrate that removing bulky surface structures improve very significantly inter-cell access to mutually matching interaction partners presented on the bacterial exterior. They also facilitate the predictive modeling of multi-strain assembly formation in tridimensional flocks embodying artificial communities or forming structured catalytic consortia.

## RESULTS AND DISCUSSION

### Identifying genetic determinants to obtain a surface-naked variant of *P. putida* KT2440

We started by an *in silico* analysis of the genome of *P. putida* KT2440 ^33^ and identified a non-exhaustive set of 21 regions/operons encoding elements that may be exposed on the bacterial surface. Within these DNA regions we included fimbriae, surface adhesion proteins, exopolysaccharides, the O-antigen side chain, the flagella, and other conspicuous envelope-associated components. These regions are distributed throughout the genome of the strain under examination (Fig. 1A). The construction of a surface-naked variant followed a specific roadmap in which the selected DNA regions were sequentially removed from the genome using a homologous recombination-based approach that generates scarless deletions ^34-35^. The specific order for the serial deletion steps followed to construct the target strain is sketched in Fig. 1B and Supplementary Table S1. Broadly speaking, we clustered these regions into four groups: [i] complex structures, [ii] surface proteins, [iii] related to production of extracellular polymeric substances (EPS) and [iv] other, non-OM associated elements.

**Figure 1.**
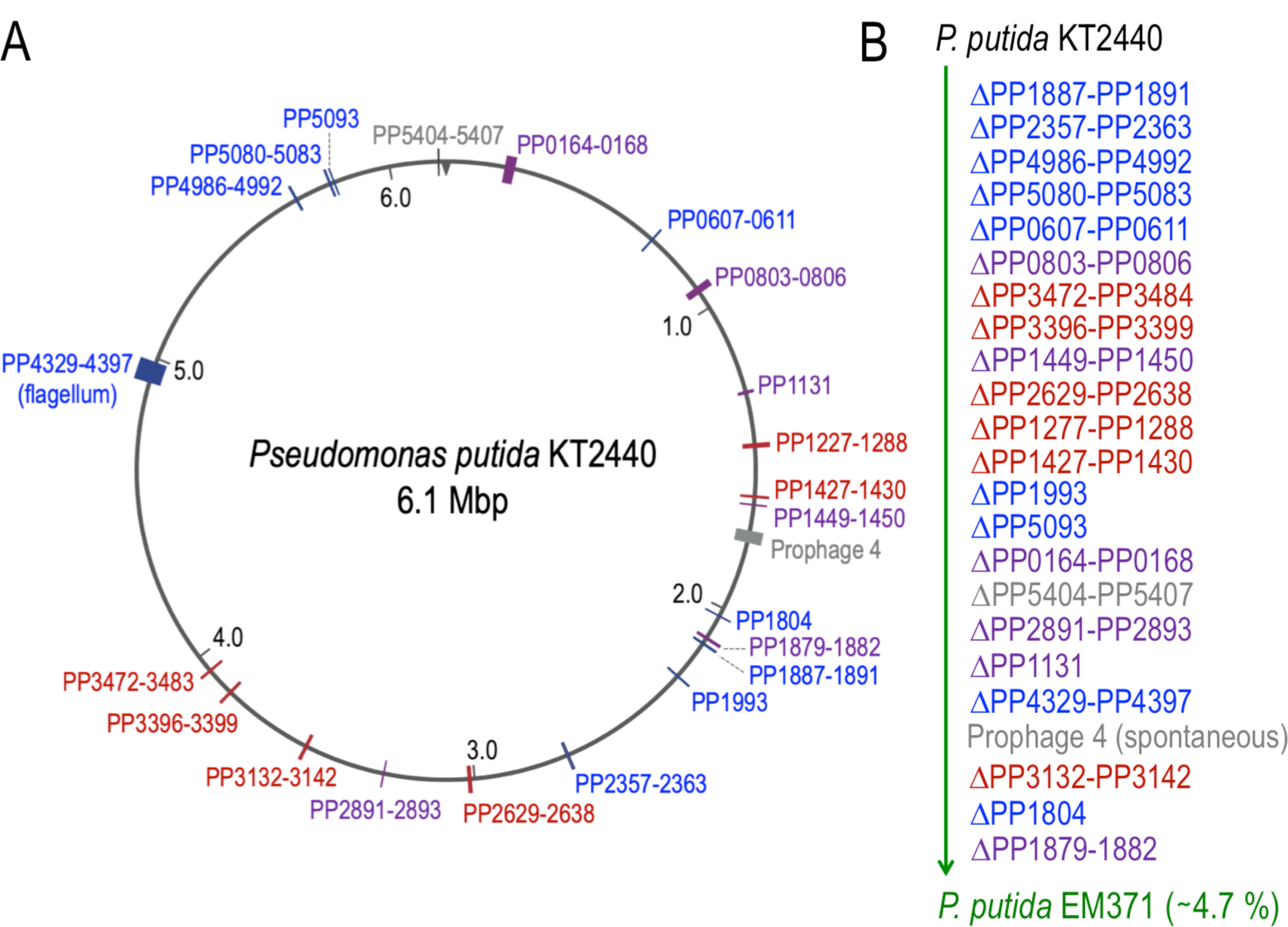
Construction of surfome-streamlined strain *P. putida* EM371. (A) genetic map of *P. putida* KT2440 depicting the approximate location of the operons deleted to engineer the surface display strain. Not drawn to scale. (B) pipeline followed to engineer the EM371 strain. The serial deletion steps with the relevant PP numbers of the regions or genes deleted are depicted in the proper order. These deletions resulted in a *P. putida* variant with a 4.7 % genome reduction. The different colors in the figure represent the artificial clustering of these regions. In blue are represented regions within the complex structures group; in purple surface proteins; in dark red the EPS-related ones and in grey the other elements.

Within the complex structures group we have two regions that encode putative fimbrial-like structures, the PP1887-PP1891 cluster and the *csu* operon (PP2357-PP2363) ^36-38^. There are also several loci that encode putative type IV pili genes and a pilus chemotaxis locus (PP0607-PP0611; PP1993; PP5080-PP5083; PP5093; and the PP4986-PP4992 respectively). These elements allow attachment to surfaces, aggregation and twitching motility ^39^. Besides, we deleted the entire flagella operon (PP4329-PP4397) ^40^. The last element of this group is the O-antigen, located at the most distal part of the complex and multidomain lipopolysaccharide (LPS) molecule. The O-antigen is composed of repeated oligosaccharides producing a side chain of variable length; it has a protective role, it is important in cell adhesion and surface attachment, and it is also highly immunogenic ^41-42^. Previous works identified that *P. putida* has only the high molecular weight B-band form ^43-44^, and in KT2440, the transmembrane glycosyltransferase WbpL (PP1804) starts the assembly of the O-antigen side chain of LPS and mutations in that gene interrupted its synthesis ^44^. Therefore, we deleted the *wbpL* gene to eliminate its synthesis, and with it also removed the last 4-bp of the upstream gene *wbpV*.

Within the second group, we selected three regions that encode large surface adhesion proteins PP0164-0168, PP0803-0806, and PP1449-PP1450 ^45^. Within the cluster PP0164-PP0168 lies the largest gene in the genome of KT2440 that encodes LapA (PP0168) a key factor for adhesion to corn seeds ^46^ and plant roots ^47^. LapA is directed to the surface by the ABC transporter composed of LapCBE (PP0166-PP0167-PP4519) ^48^. This cluster also comprises a c-di-GMP binding protein (LapD) ^49^ and a periplasmic cysteine protease (LapG) ^50^. Thus, we made a clean deletion of the whole PP0164-PP0168 cluster. We also deleted the cluster PP0803-0806 containing the second largest surface protein LapF (PP0806) and a proper ABC transporter (LapHIJ). LapF is in charge of microcolony formation by providing cell-to-cell interactions that prompt the formation of mature biofilms and it is also important in the colonization of biotic and abiotic surfaces ^51^. We also selected the operon P1449-PP1450 that encodes a two-partner secretion (TPS) system involved in seed colonization and iron uptake ^52^. Within the operon, the protein HlpA (PP1449) looks like a filamentous hemagglutinin adhesin and it is predicted to be localized either on the cell surface or secreted. Moreover, we included a putative outer membrane lipoprotein (PP1131), an operon PP2891-PP2893 encoding a putative surface antigen D15, and also the locus PP1879-PP1882 that encloses a putative outer membrane autotransporter and a large RHS element containing YD-peptide repeats.

In the EPS-related group we included exopolysaccharides and curli fibers produced by bacteria under diverse environmental conditions that have different physiological roles such as maintaining the integrity of the cell envelope, prevent desiccation, and in the colonization of surfaces by providing structural support for biofilms in many bacteria ^53^. Within the genome of KT2440 there are several loci involved in the production of different types of exopolysaccharides ^45 54^. Of these, the bacterial cellulose synthesis (*bcs*) operon (PP2629-PP2638). Cellulose is an extracellular polymer composed of D-glucose monomers linked by β-1,4 glycosidic bonds and is key for cell adherence, to provide structural support to biofilms and in the colonization of the rhizosphere ^55-56^. In a similar way, the *pea* locus (PP3132-PP3142) encodes a cellulase-sensitive polysaccharide that has a role in stabilizing biofilms ^54, 56^. Other important polysaccharide that is commonly found in the EPS matrix of bacteria is alginate, that is composed of o-acetylated β-1,4-linked D-mannuronic acid and α–L-guluronic acid ^57^. Previous reports indicated that *P. putida* produces alginate under water limitation stress in order to reduce the water loss in biofilms helping to preserve a hydrated environment and also offer protection against ROS generated due to matric stress ^58-59^. KT2440 contains two *alg* operons, one with structural genes (PP1277-1288) and another (PP1427-PP1430) with the regulatory elements and the alternative sigma factor (AlgU), that control not only alginate production but also manages cell envelope stresses ^57^. Furthermore, the proteinaceous amyloid-like fibrils known as curli play an important role in the colonization of surfaces and in the scaffolding of biofilms ^60^. The genome of KT2440 bears two clusters with components related to the synthesis and export of curli. The PP3396-PP3399 operon bears the major (CsgA-like) and minor (CsgB-like) curli subunits, where the second locus PP3472-PP3484 encodes non-structural genes involved in curli biogenesis and secretion.

Within the last group, we included an element non-related to the OM (the Tn*7*-like transposase operon PP5404-5407) in order to increase the stability of recombinant strains engineered using this mini-transposon ^61-62^. Besides, we realized that during the course of constructing this deep engineered strain prophage 4 (PP1532-PP1584) was spontaneously lost from the genome due to a natural excision event ^63^ that occurred sometime after deletion #18 (Supplementary Fig. S1A). As a consequence of that, the resulting variant is more resistant to UV than the parental strain (^63^; Supplementary Fig. S1B).

Evnetually, the surface-streamlined strain encompassed 23 deletions that include a total of 230 genes. The genomic coordinates and the precise extension of each deletion, based on the reference genome of *P. putida* KT2440 (GenBank #: AE015451), are compiled in Supplementary Table S1. Diagnostic PCRs were performed to confirm deletions using oligonucleotides that hybridize within the excised region (Supplementary Table S2 and Supplementary Fig. S1C). Also, the predicted boundaries of each deleted region were PCR amplified and the flanking regions sequenced. All of these deletions accounted for a ∼4,7% genome reduction size of the parental strain. This surface-naked variant of the wild type strain was designated as *P. putida* strain EM371. The following sections describe its phenotypic and physiological properties as well as its value for ectopic display of artificial adhesins closely bound to the cell’s exterior layer.

### The envelope-edited *P. putida* EM371 strain displays a simplified outward-facing molecular landscape

One characteristic external, bulky structure that cells expose to the outer environment is the LPS. We started by checking whether the deletion of the O-antigen glycosyl transferase (*wbpL*) correlates with the absence of this exposed structure in the *P. putida* EM371 strain. To this end, we extracted lipopolysaccharides from the wild type and the naked strain, separated them by an SDS-PAGE and visualized them by silver staining. As expected, the surface-edited bacteria lack the typical ladder pattern that correspond with the high-molecular weight of the repeating polysaccharide (Fig. 2A). This result confirms the absence of this complex structure in *P. putida* EM371. Along the same line we compared whether there were differences in the whole of the surface-exposed proteins (*surfome*, ^64-66^ detected in the parental and in the naked strain. For doing this we used an experimental approach based on the use of activated magnetic nanoparticles (NPs) combined with a multidimensional protein identification technology (MudPIT; ^67^. Briefly, small iron oxide nanoparticles (70 to 90 nm of diameter) covered with carboxymethyl dextran were chemically activated and mixed with intact whole cells as explained in the Methods section. Then, NPs attached to bacteria were magnetically purified and the bound proteins identified by MudPIT. Proteins considered bound by NPs at the cell surface were identified because the corresponding average Spectral Count (SpC) resulted significantly higher than the SpC determined with the shedding controls (see Methods for details). This technology identified a total of 196 surface-presented proteins in the parental strain while in the case of the surface-naked variant only 12 could be recognized (Fig. 2B and Supplementary Table S3). Among these, only five proteins were identified as shared by both strains. Moreover, within the proteins recognized in KT2440, those conspicuously present in the wild-type strain but missing in *P. putida* EM371 expectedly included four components of the flagellar operon and one constituent of type I fimbriae. Moreover, the pool of proteins covalently bound by NPs at the cell surface in KT2440 included typical outer membrane (OM) components such as the lipoproteins OprL and OprI, the porin OprD, OprH and an OmpA family protein (Supplementary Table S3). More surprisingly, a number of proteins predicted to localize in the inner membrane (IM) did appear as surface-exposed in the MudPIT data of Supplementary Table S3. As shown in other *surfome* studies in Gram-negative bacteria ^67^ this is not entirely unknown, as many proteins predicted to localize in the inner membrane (IM) do occasionally appear associated to the OM as well. Although technical issues cannot be ruled out, it is also possible that such proteins may reach out beyond the IM and even protrude outwards. Such a notion may also be true for a number of proteins of Supplementary Table S3, which do appear on the cell exterior despite being predicted or experimentally determined earlier within the IM. In other cases *classical* cytoplasmic proteins, such as ribosomal proteins, along with other anchorless products, have been observed on bacterial surfaces in a variety of studies ^68-71^. Data shown in Supplementary Table S3 also include some proteins typically considered cytoplasmic. Non-overlapping examples of these appeared both in the wild type strain and the surface-edited counterpart and, interestingly 7 of them i.e. PP_5354 (cell wall assembly protein), PP5001 (ATP-dependent protease ATP-binding subunit HslU), PP3431 (ThiJ/PfpI domain-containing protein), PP2410 (Cobalt-zinc-cadmium resistance protein CzcA), PP1989 (Aspartate-semialdehyde dehydrogenase), PP1360 (co-chaperonin GroES) and PP_0856 (outer membrane protein assembly factor BamB, were detected at the surface of *P. putida* EM371 and not in the parental strain. How this could happen? Specific mechanisms accounting for the observed protein misplacement cannot be ruled out, but it could also happen that alteration of the charge distribution on the cell surfaces caused by the deletions changes the profile of proteins captured by the NPs through mere electrostatic interactions. One way or the other, the drastic change in *surfome* composition and the O-antigen loss undergone by *P. putida* EM371 (Fig. 2) causes a major simplification of the molecular landscape that can be encountered when approaching cells from the outside.

**Figure 2.**
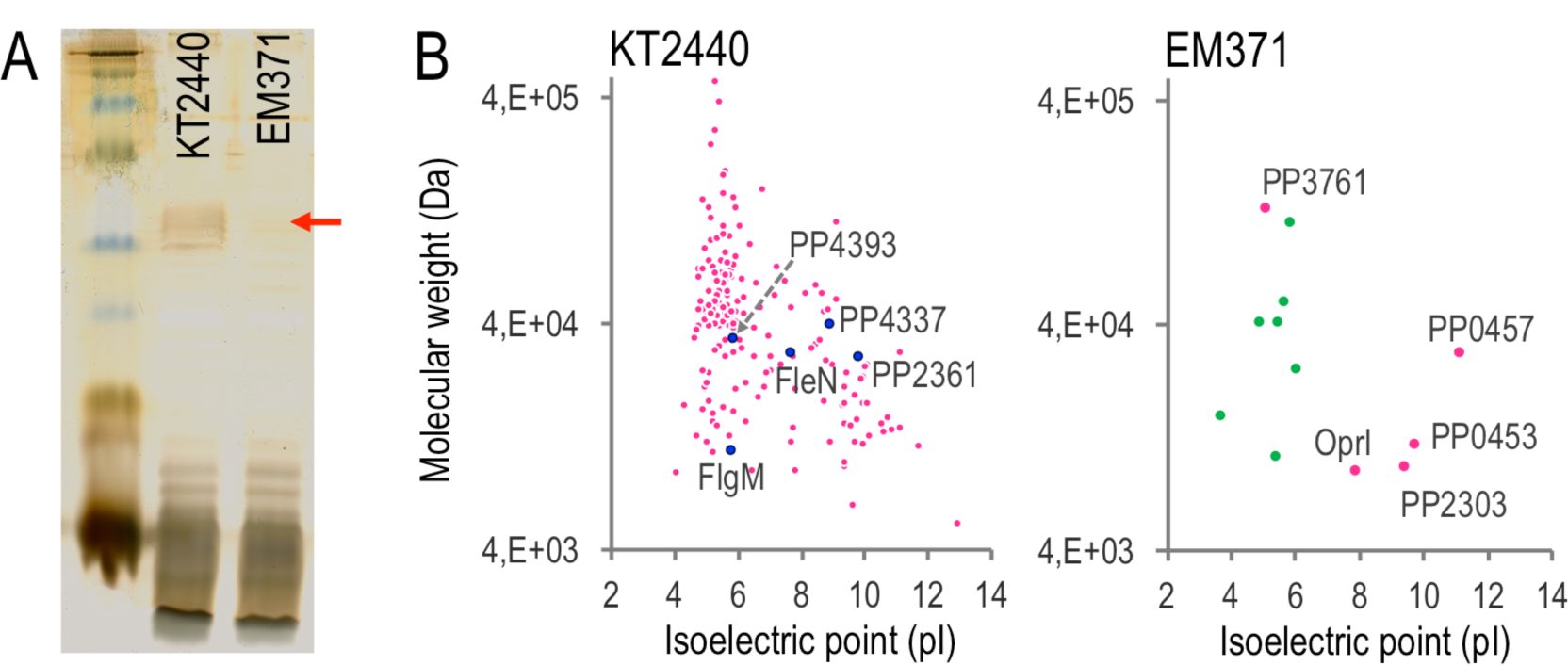
Surface properties of naked strain. (A) Silver stained SDS-PAGE to analyze the pattern of LPS in KT2440 and EM371 of whole cell lysates (represented by a red arrow in the image). Kaleidoscope pre-stained standard (BioRad, CA, USA) was used as marker. (B) surface-associated proteins captured by activated magnetic nanoparticles in *P. putida* KT2440 and EM371. Cells were treated with activated nanoparticles (NPs) to capture envelope-associated proteins that were identified through MudPIT analysis. Each identified protein is represented in a virtual 2D map by a colored point defined by its theoretical isoelectric point (pI) and molecular weight (MW). In the chart of KT2440, blue dots correspond to proteins deleted in the naked strain. In the case of EM371, pink dots correspond to five proteins also identified in the parental strain KT2440.

### Gross physiological and morphological analysis of the surface-streamlined strain

The next step in the characterization of *P. putida* EM371 was the comparison of its the physiological vigor with that of the wild type strain. This was made by assessing growth physiology on different carbon sources that establish different metabolic regimes: LB for rich media and in M9 minimal media with either a gluconeogenic (succinate) or glycolytic (glucose) carbon source. Bacterial growth was monitored by following the OD_600_ in a 96-well plate for 24h. Supplementary Fig. S2A shows the overall growth profiles of both strains with the different media under examination. There was a significant alteration in the growth rate of both strains in rich-nutrient media while there is no significant difference when growing in minimal media supplemented with either a gluconeogenic or a glycolytic carbon source. On the other hand, the total optical cell density reached with this experimental set up constraints the capacity of *P. putida* EM371 due to the combined effect of the limited aeration in a plate reader together with the absence of flagella in this strain. To clarify this we regrew cultures in Erlenmeyer flasks with vigorous shaking and measured the OD_600_ at 24 h. Under these conditions, the *P. putida* EM371 strain reached similar OD_600_ values in LB and M9 glucose while less total growth was achieved in M9 succinate (Supplementary Fig. S2B).

Since many of the elements deleted in *P. putida* EM371 may have a deep impact in the social behavior and in the cellular and macrocolony morphology of this strain, we next inspected these traits. Flagella, the locomotion organelle, is used to propel bacteria in a non-social manner to explore and scavenge nutrients in the environment ^72^. We analyzed the swimming ability of the strains of interest in soft agar plates where cells are able to penetrate the media matrix and move through the medium ^73^, thereby generating a halo that surrounds the inoculation spot. As shown in Fig 3A when cells were placed on a M9 glucose 0.3% (w/v) agar plate, deletion of the flagellar operon in the *P. putida* EM371 strain rendered non-motile cells. Morphology of individual bacteria was next examined by fluorescence microscopy. For a better visualization, cells were stained in the first case with both the membrane dye FM1-43X (that produces a strong fluorescent emission in hydrophobic environments) and DAPI (that specifically binds DNA). The differential interference contrast (DIC) and merged fluorescence images (FM1-43X + DAPI) obtained are shown in Fig. 3B. Both strains showed similar morphologies and the specific membrane dye stained both cells likewise. To obtain a closer look into the cellular morphology and also to confirm the absence of deleted outer membrane structures, we also used transmission electron microscopy (TEM) of negatively stained bacterial samples. In the pictures obtained, the flagella is clearly visible only in the wild type strain while the multi-deleted engineered strain is non-flagellated (Fig. 3C), as confirmed previously in the swimming assay.

**Figure 3.**
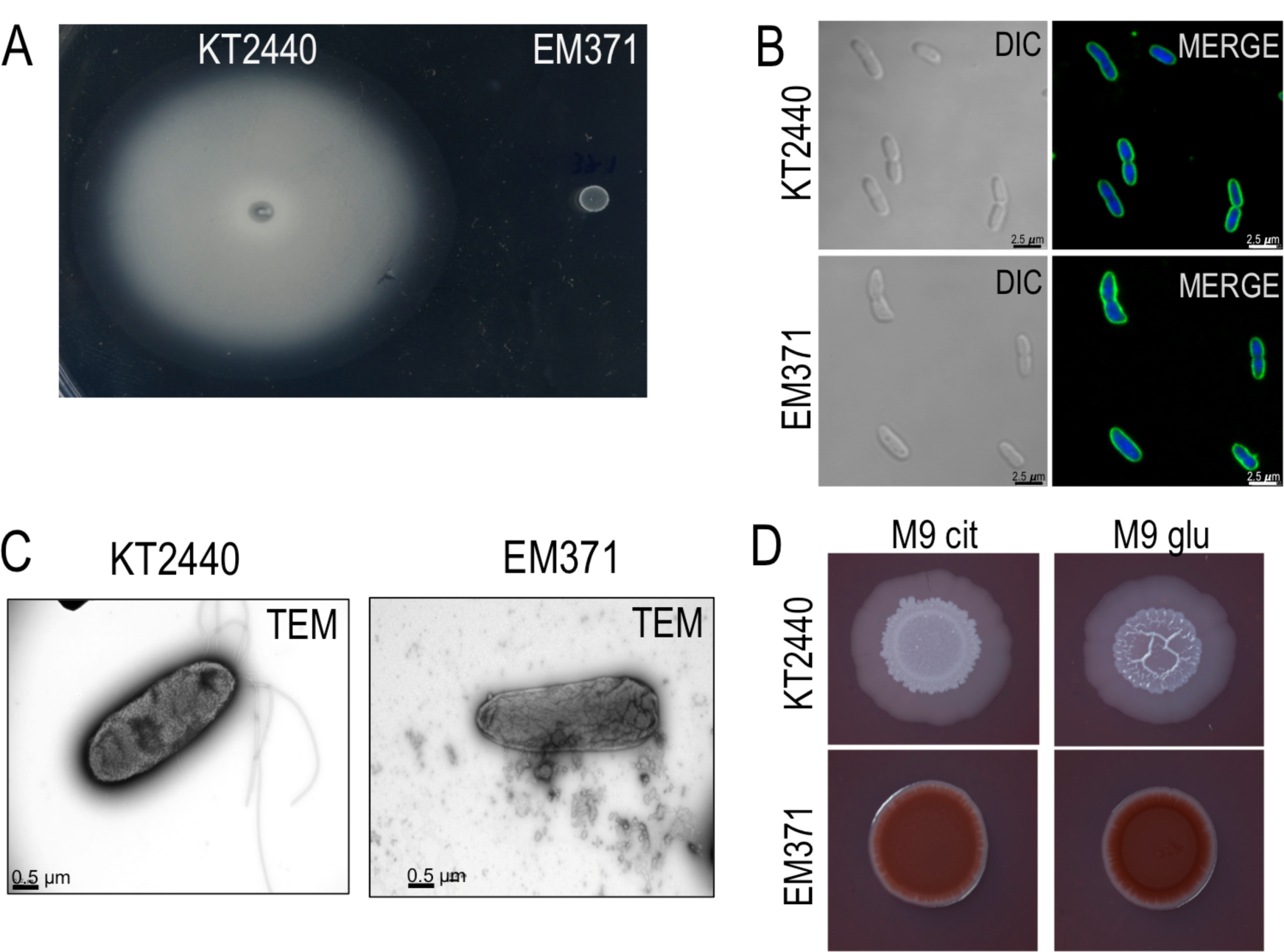
Motility test, cell and colony morphology of *P. putida* wild type and EM371 naked strain. (A) swimming ability of KT2440 and EM371. Overnight grown cells were spotted on M9-glucose semi solid agar plates and incubated at 30 °C for 48 h and photographed. (B) Nucleic acids and membrane staining of *P. putida* KT2440 and EM371. Cells were stained with the membrane dye FM1-43FX and DNA with DAPI and visualized with DIC and the appropriate fluorescence channels in a Confocal microscope. The FM1-43FX (colored in green) and DAPI (blue) channels were merged and represented. (C) electron microscopy images of the parental and naked strain. Samples were negatively stained with uranyl acetated and observed with a JEOL JEM 1011 transmission electron microscope. (D) macrocolony morphology assay. Bacterial samples were spotted on M9 minimal medium supplemented with either citrate or glucose as sole carbon source and with Congo Red and Coomassie Brilliant Blue, rested at 30 °C for 3 days and photographed.

Finally, colony morphology of the surface naked-strain was inspected as well. When bacteria grow on solid surfaces they form complex 3D multicellular structures resulting from production of a diverse number of elements commonly known as the extracellular polymeric substance (EPS), that includes exopolysaccharides, pili, adhesins and curli among others. The deletion of many of these elements in *P. putida* EM371 prompted us to explore the macrocolony development of this strain. To this end we used M9 minimal media with either citrate or glucose as C-source and Congo Red and Coomassie Brilliant Blue as generic dyes for exposing EPS production. These chemicals bind a number of extracellular matrix components such as amyloid fibers (curli), fimbriae and cellulose ^74-75^. For the assay we spotted 5 µl of overnight grown cells onto agar plates containing the dyes and incubated them for 3 days at 30 ºC. As expected the EM371 strain developed flat and smooth colonies while *P. putida* KT2440 gave rise to more complex, elaborated and wrinkled structures, especially with glucose as C-source (Fig. 3D). In contrast, *P. putida* EM371 rendered intense red colonies while the wild type strain produced much paler colors (pink to white). These differential phenotypes could be be caused by variations in the internal cyclic-di-GMP levels brought about by the resetting of the cognate regulon ^9, 76-77^). But it could also be traced to different affinities to the dyes due to altered physicochemical properties of the envelope or even to an increased self-aggregation of the engineered strain. To shed some light on these possibilities we next studied changes in the material characteristics of *P. putida* EM371’s envelope made happen by the deletion of surface structures.

### Sedimentation, surface hydrophobicity and membrane permeability

Deletion of flagella is known to influence the bacterial sedimentation rate and surface hydrophobicity of *P. putida* ^40^. On this basis, we tested how non-shaking growth conditions affects sedimentation of the engineered strain. To do that, we incubated overnight cells two grow without any agitation for 24 h at room temperature (RT) and measured the OD_600_ of the top part of the test tube. As observed in Fig. 4A, the surface-naked strain not only settled at the bottom of the tube but they did more pronouncedly that cells simply lacking the flagella ^40^. In order to determine whether this was the result of changes on cell surface hydrophobicity we next tested how the deletion of multiple outer membrane structures entered in *P. putida* EM372 strain such as flagella, bulky surface proteins, LPS etc, affected such physical parameter. For this we performed a comparative microbial adherence to hydrocarbon (MATH) test ^78-79^ with the wild type and the naked strains. This procedure measures the affinity of the bacterium to an organic phase (alkane) as a proxy of the overall hydrophobicity of the bacterial surface. The more hydrophobic the cell surface is the more bacteria partition to the organic layer and this is reflected with a higher MATH score. Not surprisingly, the naked strain showed a much less hydrophobic surface than the parental strain (*P*< 0.0001 unpaired t-test; Fig. 4B). This observation is consistent with earlier observations the effect of lacking the flagella and the LapF protein on cell surface hydrophobicity ^40, 80^ and accounts for the differential tendency to sedimentation shown in Fig. 4A.

**Figure 4.**
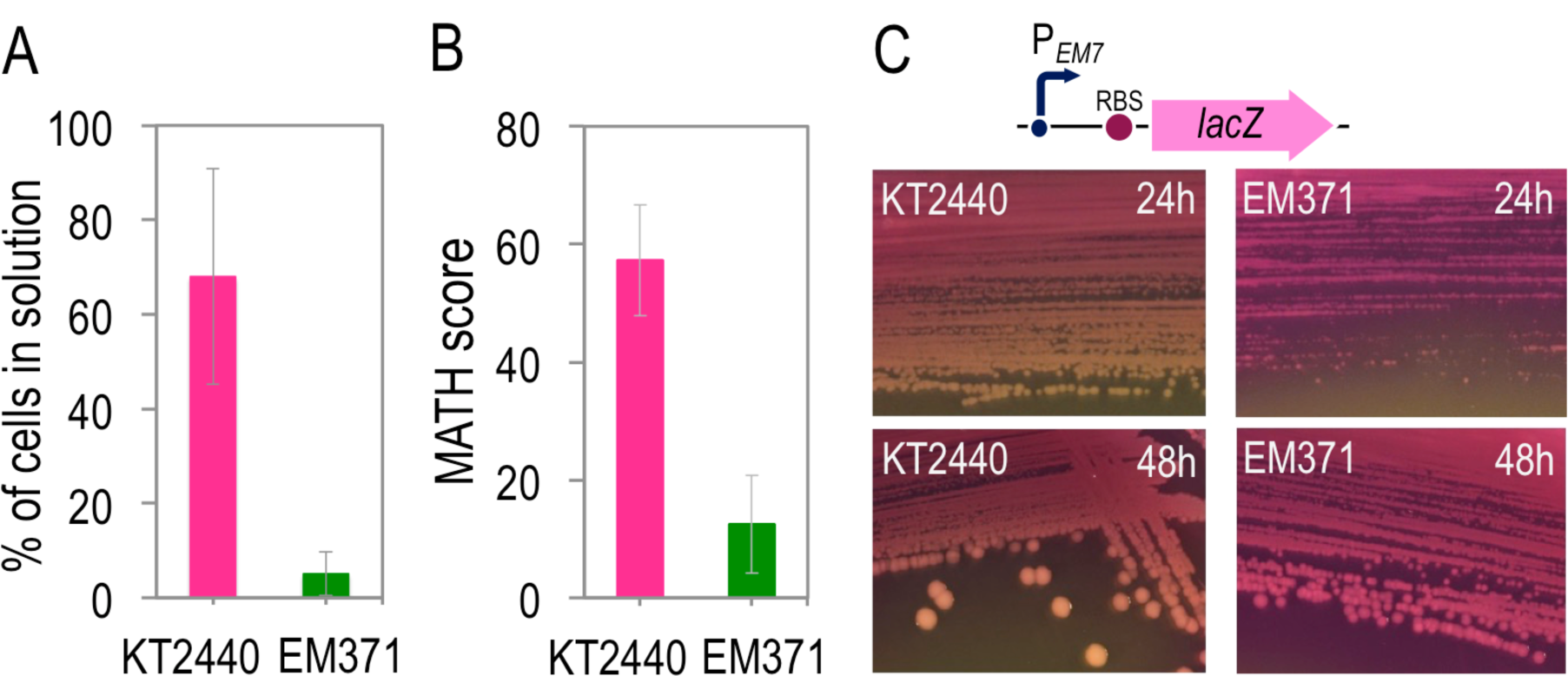
Physicochemical traits of strain *P. putida* EM371. (A) sedimentation assay of KT2440 and EM371. Overnight grown cultures in 4ml LB (OD_600_ value set to 100%) were incubated without agitation at room temperature for 24 h and the OD_600_ from the upper part of the test tube measured to calculate the percentage of cells persisting in solution for each strain. The average and standard deviation of five experiments is shown. (B) surface hydrophobicity scores for KT2440 and EM371. The microbial adhesion to hydrocarbon (MATH) experiment was performed with exponential cells washed with the PUM buffer and their adhesion to hexadecane determined to calculate the MATH score as: (1-OD_600_^final^/OD_600_^initial^)x100. The average and standard deviation of six experiments is shown. (C) permeability test by using CPRG indicator agar plates. On top, a schematic representation of the construct used to constitutively express the LacZ enzyme in the *P. putida* strains at stake. Cells, harboring plasmids with the above-mentioned operon, were streaked on LB agar plates supplemented with 20 µg ml^-1^ CPRG and incubated for 24h and 48h at 30 °C.

Deletion of the different outer membrane structures could also affect the overall permeability of the cell envelope. To check this possibility we further inspected whether the surface of the naked strain had the permeability altered using the simple and visual protocol described in ^81^. Briefly, this method relies on the use of a galactoside analog (chlorophenol red-β-D-galactopyranoside; CPRG) that produces a visible red color compound (chlorophenol red, CPR) when degraded by the cytoplasmic LacZ enzyme. Given that CPRG cannot enter cells directly, the dye becomes colored only clones with an increased surface permeability. Since *P. putida* lacks an endogenous β-galactosidase enzyme, we transformed the strains under scrutiny with high/medium copy plasmid pSEVA2513-LacZ (Supplementary Table S4), which bears the *E. coli lacZ* gene expressed from the P_*EM7*_ constitutive promoter. Transformants we then streaked on LB + Km agar plates supplemented with 20 µg ml^-1^ of CPRG, incubated and photographed after 24 h and 48 h of growth at 30°C. Fig. 4C shows that the biomass of the surface-naked strain has a pinkish-red color while parental *P. putida* KT2440 produces whiter colonies at either incubation times. This suggests that *P. putida* EM371 had a more permeable cell envelope that allows CPRG to enter cells and reach the plasmid-encoded β-galactosidase. This increased permeability may account for the red colonies that develop in media with Congo Red/Coomassie blue strain shown in Fig. 3D.

### Stress resistance profile of the *P. putida* EM371 strain

The evidence about having a more permeable envelope prompted us to test whether this was translated into increased sensitivity to archetypal stressor compounds. To examine this, we exposed cells to a number of drugs and chemicals that elicit distinct types of insults and evaluated bacterial tolerance to them by simply recording the inhibition halo caused by each specific compound (Fig. 5A). The stress response was estimated using soft-agar experiments in which a molten 0.7% (w/v) LB agar suspension containing the bacterial cells were spread onto LB agar plates. Sterile filter disks were placed on top of the bacterial lawn and soaked with the specific stressor, after which plates were incubated overnight at 30°C. Among the compounds assayed we included the β-lactam ampicillin (disrupts the cell wall by interfering with the peptidoglycan synthesis) and aminoglycosides kanamycin and gentamicin (that inhibit protein synthesis). Note that Ampicillin is a zwitteronic drug that uses porins as the conduit to cross the OM, while aminoglycosides are polycationic and hydrophobic antibiotics that gain access to the cytoplasm by permeation through the double-layered envelope ^82^. We also exposed cells to chelating agent EDTA and the denaturing anionic detergent SDS. In general, as shown in Fig. 5A, the naked strain turned out to me more sensitive to all stressors tested. The most pronounced effects were observed with the most hydrophilic ones (based on the octanol/water partition coefficient, see above), namely Km and EDTA. This confirmed that the increased permeability of the envelope of the naked strain translates into a more stress-susceptible strain. Finally, we also investigated whether high osmotic conditions and exposure to acidic and basic pH affected the viability of the surface-naked strain. In this case, cells from overnight grown cultures were serially diluted and spotted onto LB agar plates with either divergent pH values or high salt content. As observed in Fig. 5B, the surface-naked strain was also more sensitive to all of these stresses, in particular to osmotic challenge and acidic pH. Although all these tests were qualitative, taken together they expose the degree of physiological trade off between having an outer envelope more exposed to the environment and the increased vulnerability to external insults.

**Figure 5.**
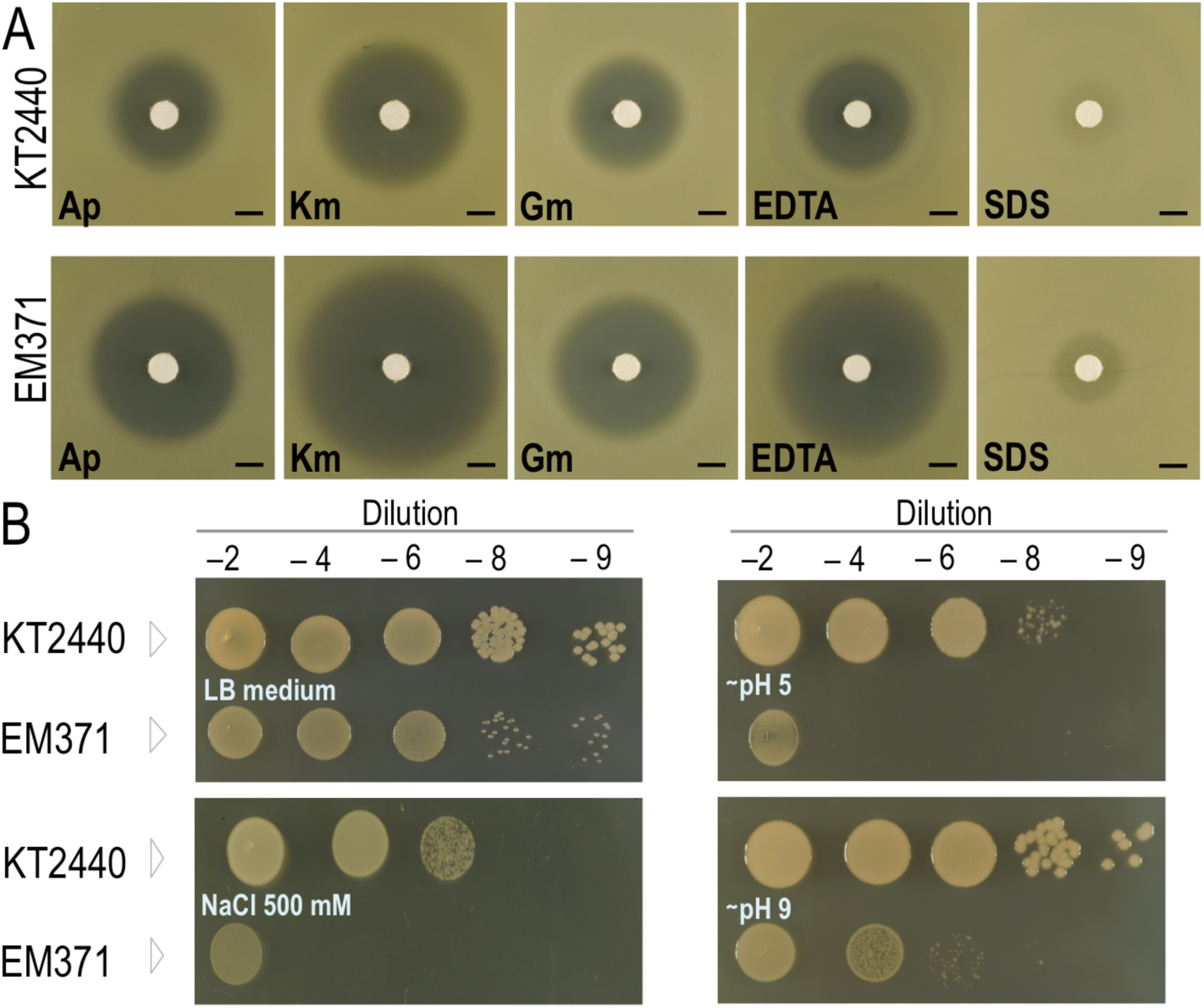
Effect of different stressors on *P. putida* KT2440 and in the engineered naked strain. (A) 100 µl of overnight grown cultures was added to soft agar and spread onto an LB agar plate. Then, a sterile filter disk spotted on the plate and soaked with 10 µl of the specific chemical (150 mg ml^-1^ Ap; 50 mg ml^-1^ Km; 10 mg ml^-1^ Gm; 0.5 M EDTA; and 10 % (w/v) SDS). Plates were incubated for 24h at 30 °C and photographed. The area that contains the inhibition halo is blown up. The black scale corresponds to the diameter of the filter disk (∼0.5 cm). (B) LB grown cultures were serially diluted and spotted onto LB agar plates supplemented with the different stressors. Plates incubated overnight at 30 °C and pictured.

### Auto-aggregation of *P. putida* EM371 depends on cell surface solvation

In view of the observations above (auto-aggregation visible with the naked-eye, alteration of surface hydrophobicity, increased permeability, sensitivity to stressors) we wondered whether this conduct could be influenced by the additives in growth media that act as solutes for solvation of the bacterial envelope. The salt (NaCl) concentration of LB medium varies among recipes from 5 g l^-1^ (85.5 mM) of LB-Lennox, to 10 g l^-1^ (171.1 mM) in LB-Miller or even to 0.5 g l^-1^ (8.5 mM) of LB-Luria. Note that the differences in osmotic pressure caused by such salt concentrations is significant. Since salt concentration does influence flocculation of bacteria in liquid cultures ^83^ we tested whether different NaCl content in the growth media modified the tendency of *P. putida* EM371 to quickly sediment. To this end, we grew the wild type and surface-naked strain in LB with concentrations of NaCl ranging 0 to 400 mM ^84^, let them grow overnight and measured the A_600_ of the liquid culture with and without vortexing. The corresponding ratio (A_600_-vortex/A_600_-no vortex) was considered have an indication of the total number of cells vs. the share of them that aggregated i.e. the bigger the ratio the more flocculation in the culture. Fig 6A shows that increasing NaCl in the growth media favored an aggregation of the surface-naked bacteria. In contrast, such salt-dependent sedimentation was not observed in the parental strain. Since solvation of the cell surface allows particles/bacteria to remain suspended in liquid media, this phenomenon is likely to reflect the cumulative result of the absence of the O-antigen and other exposed proteins in the interactions of charged sites of the envelope with salt’s ions. Such surface structures contribute to solvate the cell envelope of the parental strain. In contrast, the observed auto-aggregation of *P. putida* EM371 plausibly reflects a phase separation phenomenon similar to the salting-out effect of proteins ^85^.

**Figure 6.**
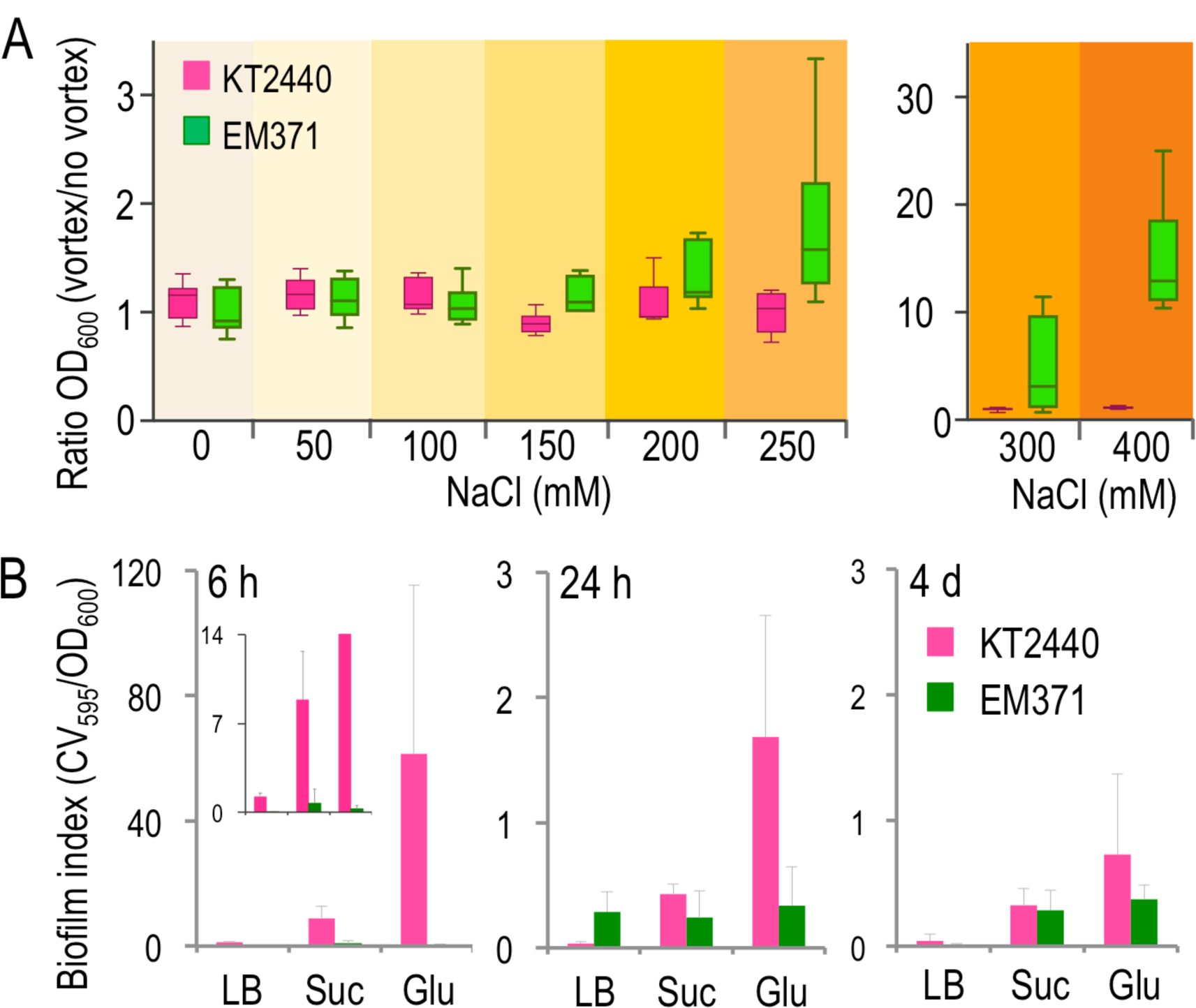
Aggregation and biofilm formation. (A) Influence of NaCl in the growth physiology of KT2440 and EM371. Cells were grown on LB without and with different NaCl concentrations overnight with 170 rpm at 30 °C. Then, the OD_600_ of cultures measured before and after vortexed the tubes to disrupt flocks. The OD_600-_vortex was divided by the OD_600_-no vortex and the obtained values of six experiments are represented as a Tukey box and whiskers plot. Pink color is for KT2440 while green is for EM371. (B) biofilm index after 6h, 24h and 4 days of KT2440 (pink) and EM371 (green) in different media: LB as rich media, M9 minimal media with either 0.2 % (w/v) succinate (Suc) or glucose (Glu) as C-source. Biofilm index was calculated as the ratio of crystal violet staining (CV_595_) to the OD_600_ of the culture. Within the 6h chart, a zoom up has been included to better visualize the biofilm response of both strains at lower values. The average and standard deviation of three experiments are shown.

In this case, particles of amphiphilic behavior are segregated from the solvent by increasing the ionic strength of the medium. Under this scenario, increasing NaCl concentration fosters cell clumping by hydrophobic interactions, as the solvating power of the surface of the *P. putida* EM371 strain would not be strong enough to thermodynamically stabilize cells in solution, and counteract this effect ^86-88^. The practical consequence of all this is that surface-naked cells can be made more or less stable in suspension by manipulating salt contents of the growth media.

### Attachment of surface-naked bacteria to solid supports

The next step in the characterization of *P. putida* EM371 involved inspection of whether this strain attached to abiotic surfaces and could establish biofilms. Since the surface-naked variant lacks elements that are crucial for bacterial adhesion and biofilm development we expected a reduction of this capacity in comparison with the parental strain. To test this, we analyzed biofilm formation at different physiological states and with various growth media to cover a broad number of conditions. Biofilm formation was analyzed through the classical crystal violet assay ^89^ using three different time points that represent the initial (6h), middle (24h) and latter (4 days) stages of the process. We also compared biofilm formation in three different growth conditions: rich media (LB), gluconeogenic (M9+succinate) and glycolytic (M9+glucose). As shown in Fig. 6, the surface-naked strain did not form as much apparent biofilm as the wild type in any of the conditions tested, specially at the initial stages of biofilm formation (6h). At this time, flagella initiates the attachment process followed by the action of pili, surface proteins and EPS to complete the formation of the biofilm structure ^89-90^. At the 24 h time point, the naked strain produced less biofilm than the wild-type bacteria in minimal media. In contrast, *P. putida* EM371 remained stably stuck to solid surfaces in LB medium. This phenomenon could be due to the flagella-mediated disengagement of *P. putida* KT2440 cells from matured biofilm to colonize new niches ^91-92^. In this case, the surface-naked cells could not escape from their initial landing pad, thereby remaining in a sessile form. The tendency remains after 4 days: by that time the wild-type cells have gone through all phases of biofilm development, while the surface-naked cells display a low capacity of adhesion to the wells throughout the whole timeline of the experiment. In fact, it is very unlikely that *P. putida* EM371 forms *bona fide* biofilms at all: what we see as crystal violet stain-able biomass is likely to be an amorphous aggregate rather than a typically 3D-structured bacterial population. Once these qualities of the surface-naked strain were set we moved on to the ultimate tests for assessing accessibility of the outward-facing cell envelope to exterior actors.

### Displaying heterologous autotransporters with Fos and Jun interacting partners in *P. putida* EM371

As mentioned above, one key motivation for constructing a surface-naked strain was to facilitate the display of molecules on the bacterial body and thus modify adhesion properties *a la carte*. To test the value of the *P. putida* EM371 strain to this end we recreated in *P. putida* the protein display system described for *E. coli* by ^11^. This approach consists on the production of a hybrid polypeptide containing the β-domain of the immunoglobulin A protease from *Neisseria gonorrhoeae* (IgAβ) fused to the eukaryotic leucine zipper transcriptional factors Fos or Jun. In that way, the IgAβ crosses the inner membrane thanks to the presence of the *pelB* signal peptide (SP) at the N-terminal of the hybrid protein and the β-domain integrates in the outer-membrane, thereby exposing the Fos or Jun molecules to the external medium. These leucine zippers tend to interact through the coiled coil domains of their structure and produce heterodimers (also the Jun molecules could interact with each other producing homodimers ^93^). On this basis, we cloned the Fos-E-tag-*igA*β and the Jun-E-tag-*igA*β modules into the broad host range expression vector pSEVA238 to obtain plasmids pSEVA238-AT-Fos and pSEVA238-Jun (Supplementary Table S4). As an intracellular expression control, we cloned the hybrid nanobody Trx-^G6^V_HH_ from ^94^, into the same backbone to yield plasmid pSEVA238-trx-^G6^V_HH_ (Supplementary Table S4). A schematic representation of the business parts of these three plasmids is shown in Fig. 7A. They were then introduced into the fluorescently-tagged variants of the naked strain (non-labelled, GFP and mCherry; Supplementary Table S4) and expression of hybrid proteins Fos-*igA*, Jun-*igA* and Trx-^G6^V_HH_ was induced upon addition of inducer 3-methyl-benzoate (3MBz) to the growth media. In vivo expression and cellular localization was then monitored owing to the E-tag epitope engineered in the fusions (Fig. 7A). Fig. 7B shows the Western blot corresponding to whole lysate of induced cells of the surface-naked strain treated (+) or not (-) with a protease (trypsin). This enzyme cannot penetrate intact cells and thus degrades only exposed and accessible proteins ^95-96^. The Western blot result of Fig. 7B shows that the hybrid Fos-*igA* and Jun-*igA* are expressed in the surface-naked strain (∼54 kDa) and that both are clearly located on the cell exterior since those bands disappeared after the treatment with trypsin. In the case of cells bearing the control plasmid with an intracellular protein, expression (∼29.5 kDa) is observed independently or not of trypsin treatment. Supplementary Fig. S3A shows that the cytoplasmic control construct (Trx-G^6^V_HH_) is not degraded by the protease unless cells are made permeable with lysozyme. Note that the effect of the enzyme was verified by immunofluorescence (Supplementary Fig. S3B). When the same plasmids were placed in the parental strain *P. putida* KT2440, correct expression and surface localization of Fos-*igA*, Jun-*igA* and Trx-^G6^V_HH_ could be verified as well (Supplementary Fig. S4). However, this data do not tell us by themselves how easily the exposed Fos and Jun moieties of the hybrid proteins are accessible from the cell’s outside in each strain. To test this, we resorted to a simple test for detecting surface-to-surface contacts as explained below.

**Figure 7.**
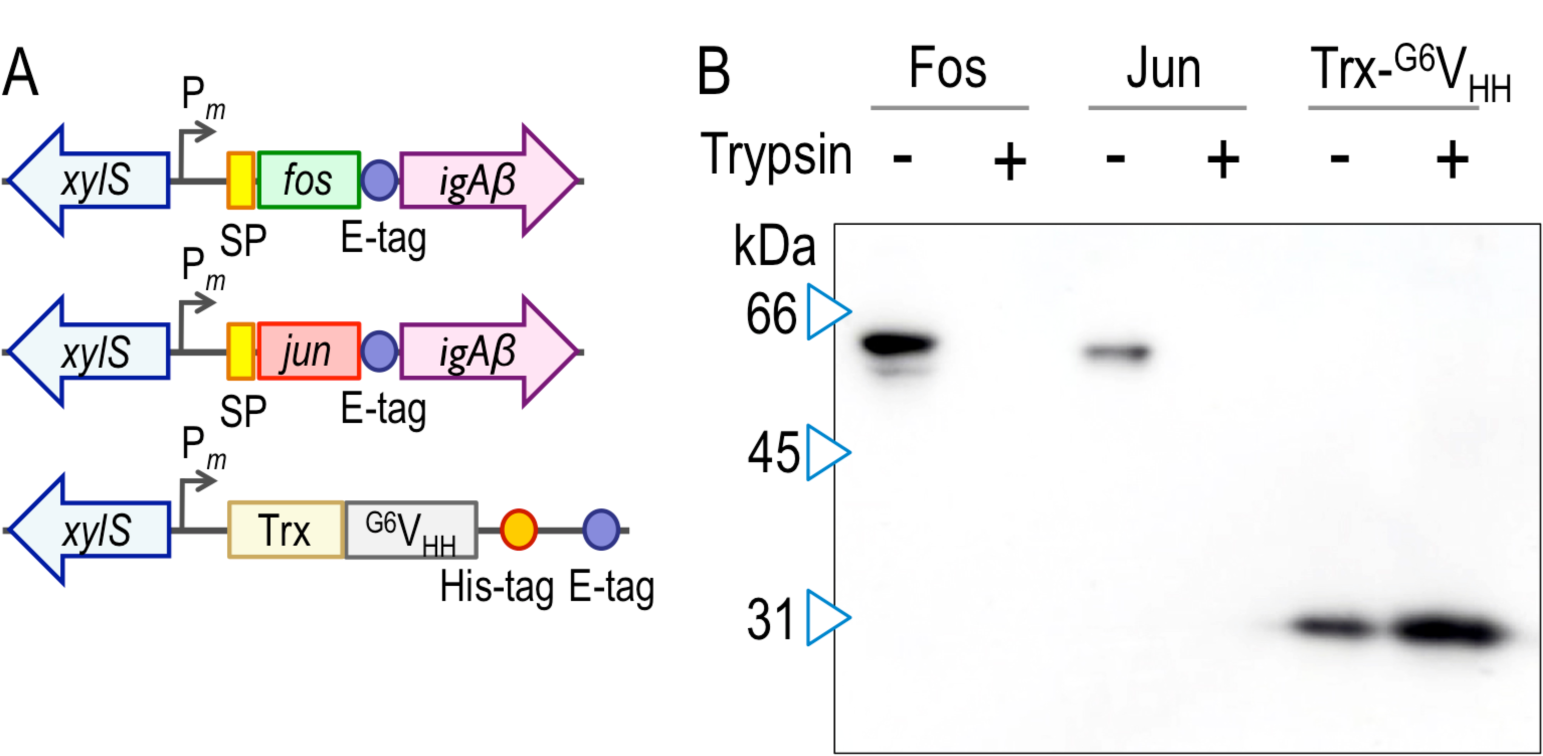
Organization, expression and localization of the Fos and Jun chimaeric proteins. (A) schematic representation of the constructs used for surface display in *P. putida*. The fos and jun leucine zipper domains preceded by the *pelB* signal peptide sequence (SP; yellow) and together with an E-tag epitope (blue circle) and the *igA*β module as the transporter module (purple) were placed under control of the *xylS*-P_m_ expression system of pSEVA238 ^27^. As a cytoplasmic expression control, we used a construction cloned into the same expression pSEVA238 plasmid containing a thioredoxin domain (Trx) fused to the ^G6^V_HH_ nanobody followed by the His-tag and E-tag epitopes. (B) western blot of induced whole cell extracts of EM371 with pSEA238 plasmids containing either the Fos-*igA*β, Jun-*igA*β or the Trx-^G6^V_HH_ hybrid proteins. Also, induced cells were treated (+) or not (-) with trypsin. The western blot was revealed with anti-E-tag as the primary antibody and reveled with anti-mouse IgG conjugated with peroxidase.

### Conditional flocculation of *P. putida* upon expression of surface-displayed synthetic adhesins

To have an estimation of accessibility of *P. putida* surfaces under various conditions, we developed the aggregation test shown in Fig. 8A. Green-tagged and red-tagged *P. putida* cells expressing Fos-E-tag-*igA*β and Jun-E-tag-*igA*β were mixed in a 1:1 ratio and production of the corresponding proteins induced with 3MBz. The rationale of the experiment is that display of that matching interacting partners should bring about cell-to-cell specific interactions that grossly manifest as aggregates of different sizes within the culture. As shown in Supplementary Fig. S5, the aggregation caused by expression of the artificial adhesins (F-J and J-J) in *P. putida* EM371 became evident when inspected in detail with confocal microscopy.

**Figure 8.**
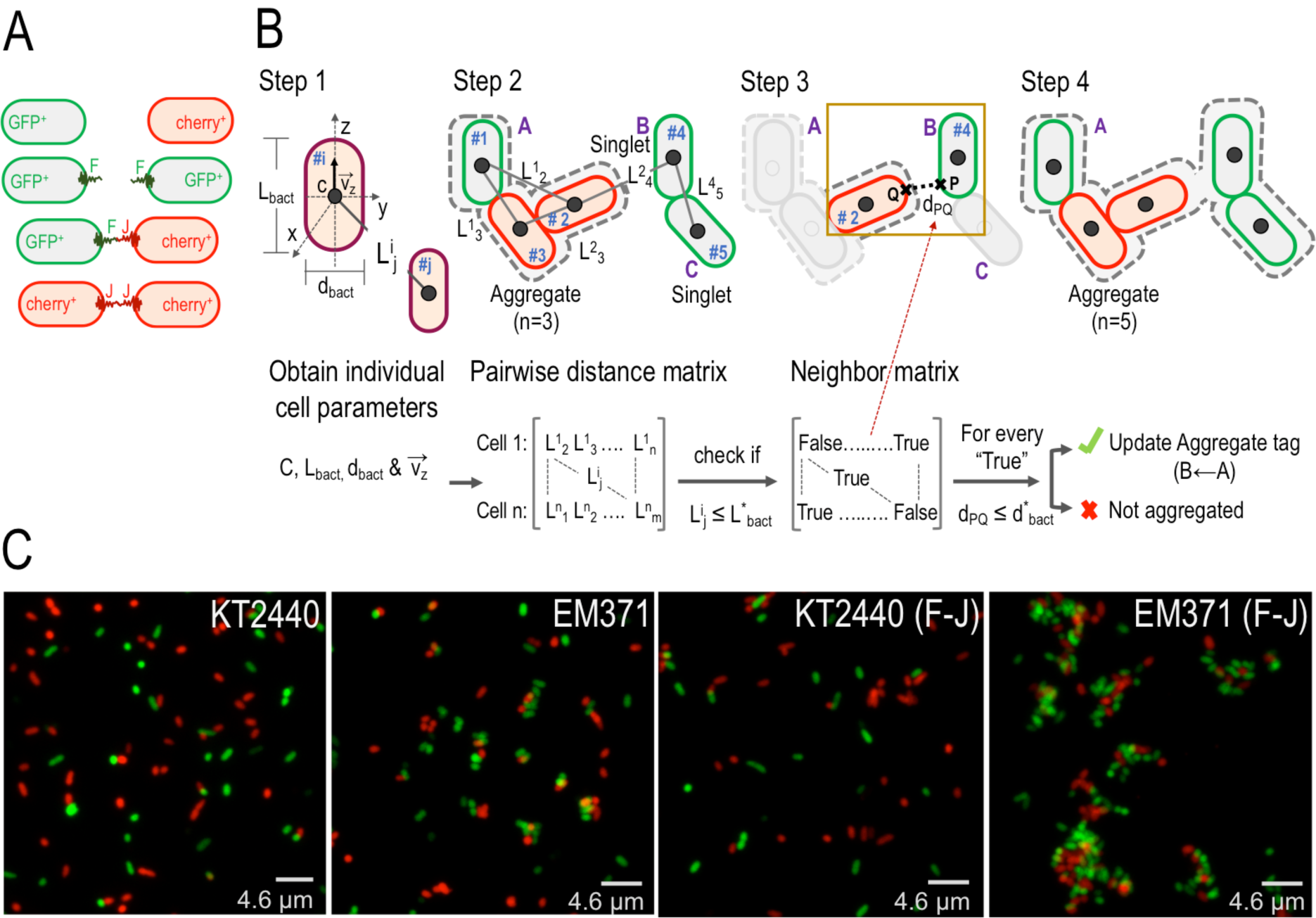
Modelling and visualizing *P. putida* cell-cell adhesion. (A) predicted possible bacterial pair interactions between strains expressing either Fos and Jun or Jun and Jun. KT2440 and the EM371 strains were fluorescently tagged with GFP and mCherry. GFP-labelled strains were transformed with Fos expressing plasmids (pSEVA238-AT-Fos) while cherry cells with Jun (pSEVA238-AT-Jun). (B) Steps followed by the in house designed computational workflow to identify aggregate clusters within confocal microscopy images. Basically, the algorithm follows 4 steps. Step1, gathered microscopy images were analyzed using Imaris software to obtain geometrical parameters of individual cells such as the mass center position (c), length (L_bact_), diameter (d_bact_) and the axial orientation vector (**v**_**z**_). Then, in step 2 distances between cell pairs were computational arranged into a distance matrix (L^i^_j_ represents the distance between the bacterial centers of #i to #j), where each row contains all distance pair combinations of one bacterium (eg: #1) to the rest of cells indicated (L^1^_2_, L^1^_3_… L^1^_n_). After that, in step 3, it was checked that those pair distances (L^i^_j_) were smaller or equal than a threshold distance (L^*^_bact_; calculated as the L_bact_ average of all cells of all images) to detect potential contacts. Such logic evaluation generates a Boolean neighbor matrix were distances satisfying the previous criterium are labelled as “true” and “false” otherwise. Finally, in step 4, every potential contact is confirmed by computing the shortest distance between those cell pairs (d_PQ_) by using a second proximity criterium (d_PQ_ ≤ d^*^_bact_; being d^*^_bact_ the average of all cell diameters detected in all images). (C) representative confocal images of the four types of aggregation experiments performed. In this, GFP and mCherry labelled strains without plasmids (KT2440 and EM371) or with the appropriate pSEVA238-AT-Fos and pSEVA238-AT-Jun plasmids were mixed at 1:1 ratio and imaged.

In order to parameterize the effect of the surface editing made in the *naked* strain we developed an algorithm for inspecting quantitatively microscope images, including physical identification and counting of either bacterial aggregates (n > 2; in order to exclude possible diving cells) or singlets (n ≤ 2) in each condition (Fig. 8B). Aggregation data were then generated by inducing cultures containing equal amounts of either *P. putida* EM371-GFP (Fos-*igA*) or *P. putida* KT2440-GFP (Fos-*igA*) with either *P. putida* EM371-cherry (Jun-*igA*) or *P. putida* KT2440-cherry (Jun-*igA*). Samples were then observed under confocal microscopy and the percentage of cells forming aggregates versus singlets automatically quantified. Fig 8C shows representative pictures of strains *P. putida* EM371 and *P. putida* KT2440 with the Fos and Jun expressing plasmids as well as same strains without the adhesins. The resulting quantification indicated that 84% of *P. putida* EM371 cells expressing Fos-*igA* and Jun-*igA* formed *bona fide* aggregates, while those formed by adhesin-less bacteria went down to 54%. In the case of the wild-type strain *P. putida* KT2440, the percentage of aggregates mediated by adhesins was 27%, while spontaneous clumpling came down to 9% (Fig. 9A). Furthermore, *P. putida* EM371 cells expressing adhesins resulted in aggregates containing more bacteria in average (n = 5) than in any of the other cases (Fig. 9B and Supplementary Fig. S6). Despite the above-discussed tendency of *P. putida* EM371 strain to flocculate naturally, expression of artificial adhesins enhanced cellular aggregation by a factor ≥1.8 (Fig. 9C). On this basis, we estimated that external approachability of the cell surface nearly doubled after stripping the bacterial envelope of all structures listed in Fig. 1.

**Figure 9.**
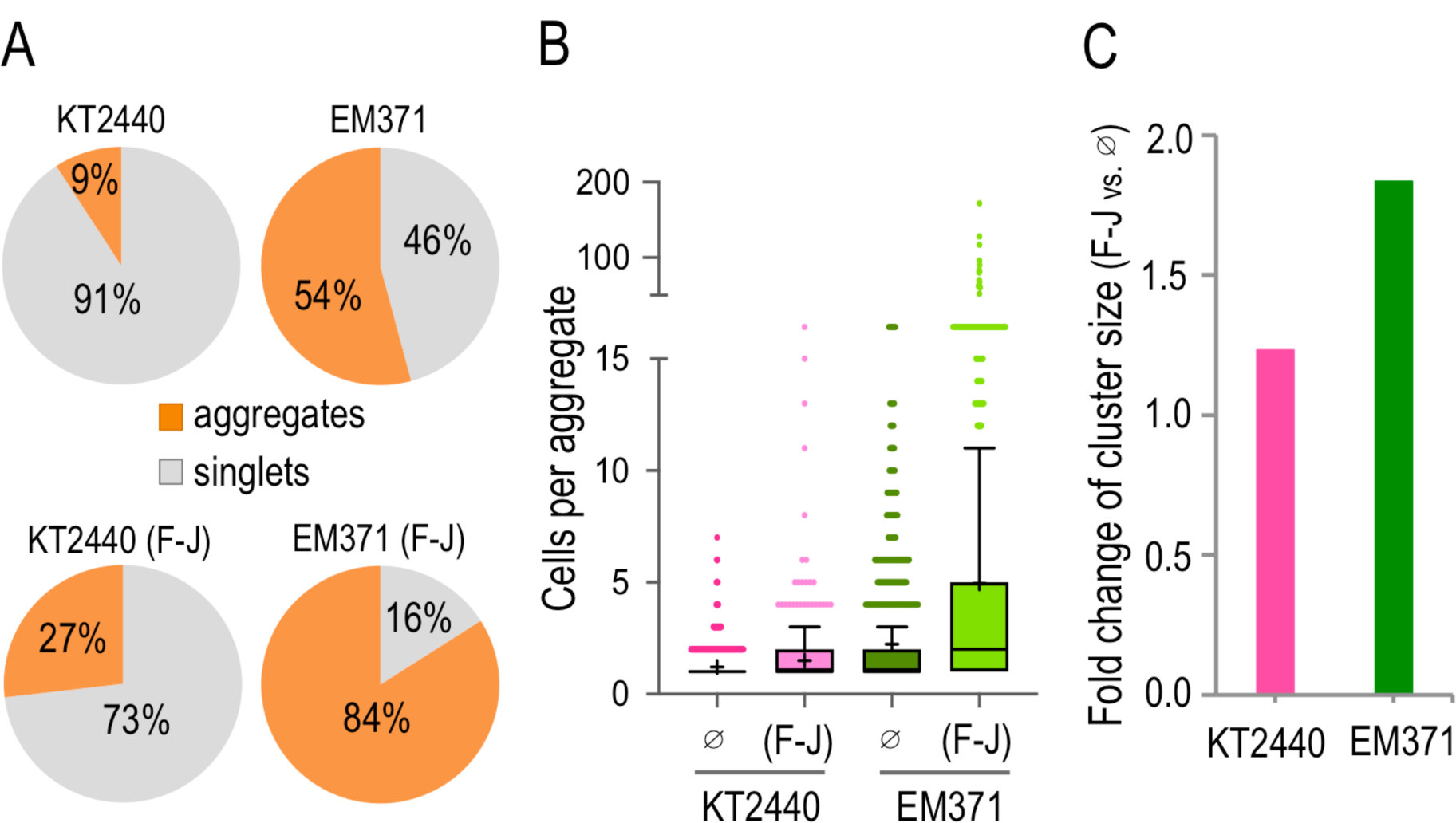
Parameterization of adhesin-mediated *P. putida* cell-cell interplay. (A) Pie charts showing the percentage of cells found as singlets (number of cells ≤ 2) versus aggregates (n > 2) in the images processed. (B) number of cells per aggregate in each condition represented as a Tukey box and whiskers plot. The average is represented within a cross while the median with a horizontal line. Values plotted outside the box represent outliers; meaning that their value is 1.5 times the interquartile range (IQR), either below (Q1-1.5xIQR) or above (Q3+1.5xIQR). (C) fold change of cluster size in both strains. This parameter was calculated as the ratio of the average cluster size in the F-J condition to the ones found in the strain without plasmids (∅). The average cluster size was estimated as the fraction of cells identified in aggregates divided by the number of clusters. *P. putida* KT2440 is represented with a pink bar while EM371 with a green one. For this experiment, two biological replicates and a minimum of 32 images were analyzed.

### Conclusion

While the value of *P. putida* KT2440 and its derivatives as a platform for metabolic engineering and a Synthetic Biology chassis is well accredited ^24, 31^, there is still much room for improving its performance through engineering physical—not just biochemical—properties of the bacterial cells. One appealing scenario deals with ectopic display of enzymatic activities or other functional structures on the cell surface ^97-99^. Moreover, rational design of single-strain and multiple-strain consortia with a given composition and 3D architecture is bound to boost the industrial and environmental uses of SynBio agents ^2, 20, 100^. Yet, naturally-occurring bacteria are most often set evolutionarily with traits at odds with such engineering objectives. In the specific case addressed in this article, the complex organization of the *P. putida*’s cell surface not only diverts a good share of metabolic resources for deployment of structures (e.g. EPS, LPS, flagella, fimbria etc) that are not only useless in a reactor (or biotechnological setting thereof) but it also limits the spatial accessibility to new functions engineered on the bacterial surface. The results shown above expose the value of stripping the envelope of *P. putida* KT2440 of a number of bulky components to increase exposure of the OM surface to the external medium. The result of this endeavor was *P. putida* EM371, which lost some typical characteristics of strains of this genus but also gained a superior ability to present heterologous proteins anchored on the cell body to the external medium. In this work, this was verified by inspecting surface accessibility of Leu zippers, which barely protrude the OM by a few åmstrongs upon display with an autotransporter protein carrier ^101^. The utility of the strain to this end has been recently exploited for functional display other, much bigger structures e.g. cellulosome scaffoldings ^102^. The superior ability of *P. putida* EM371 to ectopically display such proteins is the starting point for a series of strains specialized on surface display of growingly complex structures.

## METHODS

### Bacterial strains, plasmids, culture media and growth conditions

The bacterial strains and plasmids used in this work are shown Supplementary Table S4. As rich medium we used LB (10 g l^-1^ tryptone, 5 g l^-1^ yeast extract, and 5 g l^-1^ NaCl) and for minimal media we used M9 ^103^ supplemented with different carbon sources at a final concentration of 0.2 % (w/v). *P. putida* was incubated at 30 ºC while *E. coli* cells were grown at 37 ºC. Antibiotics were used at the following final concentrations: 150 µg ml^-1^ ampicillin (Ap) for *E. coli* and 500 µg ml^-1^ for *P. putida*; 50 µg ml^-1^ kanamycin (Km). To select appropriate clones with the alpha complementation procedure we added 40 µg ml^-1^ 5-bromo-4-chloro-3-indolyl-β-D-galactopyranoside (X-gal), plus 1.0 mM isopropyl-β-D-1-thiogalactopyranoside (IPTG) to the LB agar plates. The growth kinetics of both strains with different nutrient conditions was obtained by following the optical density (OD) at 600 nm of the cultures, inoculated at an OD_600_ of 0.05, in 96-well microtiter plates using a SpectraMax M2^e^ microplate reader (Molecular Devices, Sunnyvale, CA, USA). In the case of the final OD_600_ at 24 h, cells of both strains were grown overnight in the appropriate media, then diluted to obtain an initial OD_600_ of 0.005 in 10 ml of the corresponding culture medium and grown at 30 ºC with agitation (170 rpm) in 50-ml Erlenmeyer flasks. Then, cultures were vortexed to eliminate clumps and the OD_600_ measured after 24 h of growth.

### General DNA techniques

DNA was manipulated using common laboratory techniques described in ^103^. Plasmid DNA was prepared using the QIAprep Spin Miniprep kit (Qiagen, Inc., Valencia, CA). When required DNA was purified using the NucleoSpin Extract II (Macherey-Nagel, Düren, Germany). The oligonucleotides used in this work are indicated in Supplementary Table S2 in the Supplementary Material. Colony PCR was performed by transferring cells with a sterile toothpick directly from fresh agar plates into PCR reaction tubes. Suitability of constructs was confirmed by DNA sequencing using primers described in Supplementary Table S2. Fluorescent derivative strains were obtained by site-specific integration of mini-Tn*7* derivatives bearing constitutively expressed either GFP or mCherry fluorescent reporter cassettes ^104-105^.

### Genome deletions

Genome deletions were performed using the I-*Sce*I protocol described in ^34-35^. Concisely, upstream (TS1) and downstream (TS2) regions of homology were PCR amplified and both TS1 and TS2 fragments glued together by means of SOEing PCR ^106^. The subsequent TS1-TS2 DNA fragment was digested with appropriate enzymes and cloned into the linearized pEMG plasmid. After that, recombinant plasmids were transformed into *P. putida* cells bearing the pSW-I plasmid, ^107^ to obtain cointegrates. Once cointegrated clones were obtained, the second recombination event was forced by inducing the expression of the I-*Sce*I endonuclease ^35^. Then, Km-sensitive clones were checked by PCR to confirm the corresponding DNA deletion. Finally, after the last genome deletion was done, the pSW-I plasmid was eliminated by growing cells without selective pressure, and the plasmid loss confirmed first by sensitivity to Ap (500 µg ml^-1^) and then with colony PCR using oligonucleotides that hybridized with the plasmid backbone (Supplementary Table S2 in the Supplementary Material).

### Construction of plasmids for surface display in *P. putida*

Plasmids pJun*β* and pFos*β*, containing the β-domain of the *igA* protease fused to the transcriptional leucine zippers Jun and Fos were obtained from ^11^. Those constructs were digested with the restriction enzyme NotI and the 1.6-kb fragment cloned into the pSEVA238 expression plasmid ^108-109^. The plasmids generated were named pSEVA238-AT-Jun and pSEVA238-AT-Fos (Supplementary Table S4). The negative control plasmid pSEVA238-trx-^G6^V_HH_ was constructed as follow: pSEVA238-trx-^G6^V_HH_ was generated upon digestion of pSEVA-T•G6^ATG^ plasmid ^94^ with SacI/XbaI and the 0.8 kb resulting fragment ligated into the pSEVA238 vector.

### Morphological and phenotypic assays

For fluorescence microscopy, overnight cultures were diluted in phosphate buffered saline (PBS; 8 mM of Na_2_HPO_4_, 1.5 mM of KH_2_PO_4_, 3 mM of KCl and 137 mM of NaCl, pH7) and stained with 5 µg of FM1-43FX for 10 minutes at room temperature. Next, cells were washed and resuspended in 1 ml of PBS. Then, a 3 µl of the membrane-stained cells was transferred onto an air-dried coverslip coated with 0.1 % (w/v) poly-L-lysine. Finally, DNA was stained with 100 µl of DAPI (4’,6’-diamidino-2-phenylindole; 2 ng µl^-1^) for 2 minutes. Images were taken with a Confocal multispectral Leica TCS SP5 system (HCX PL Apo CS 100 × 1.4 oil). For quick inspection of bacterial aggregates, overnight grown cultures were diluted to an OD_600_ of 0.1 and grown at 30 ºC until OD_600_ 0.5. At that point, cultures were induced with 1mM 3MB and let them grow for 3h at 30 ºC. After that, samples were mixed at 1:1 ratio in 10 ml test tubes and let them stand for 20 minutes at room temperature. Then, 5 µl of the bacterial culture placed onto a slide with 5 µl of Prolong (Life technologies; Thermo Fisher Scientific) and covered with a 0.1 % (w/v) poly-L-lysine coverslip. Samples were visualized in a Leica DMI600 B fluorescence microscope. To prepare samples for electron microscopy, cells were laid onto carbon-coated collodion grids, negatively stained with 1 % (w/v) uranyl acetate, and samples observed with a JEOL JEOM 1011 transmission electron microscope. To quickly evaluate the lack of motility in the naked strain, 2 µl of overnight cells spotted onto M9 minimal media with 0.2 % (w/v) glucose and solidified with 0.3 % (w/v) agar, and plates incubated at 30 ºC for 48h. Bacterial colony morphology was inspected by spotting 5 µl of overnight grown cultures on M9 minimal media with 0.2 % (w/v) glucose or citrate as carbon source with 40 µg ml^-1^ Congo red and 20 µg ml^-1^ Coomassie Brilliant Blue. Plates were incubated at 30 ºC for 3 days and photographed with a Nikon D60 equipped with an AF-S Micro Nikkor 60 mm f/2.8G ED. For the analysis of the O-antigen we followed the protocol described in ^43, 110^. Briefly, cells were grown overnight aerobically at 30 ºC, then the OD_600_ of cultures adjusted to 0.4, washed twice with PBS, lysed in SDS-PAGE sample buffer and treated with proteinase K at 60 ºC for 1 hour. After that, lipopolysaccharides were separated in a 15 % (w/v) SDS-PAGE and visualized by silver staining. We used the MATH score as a proxy to assess surface hydrophobicity ^78-79^. For that, exponential cells (OD_600_ ∼0.6) were washed with phosphate-urea-magnesium sulphate buffer (97 mM K_2_HPO_4_, 53.3 mM KH_2_PO_4_, 30 mM urea, 815 µM MgSO_4_; pH7.1). Then, the OD_600_^initial^ measured and 1.2 ml of that cell suspension mixed with 0.2 ml of hexadecane, vortexed for 45 seconds, incubated stand for 30 minutes at RT and the OD_600_^final^ of the aqueous phase measured; thus the MATH score calculated as: 1-OD_600_^final^/OD_600_^initial^) x 100. The sedimentation of strains was observed as follows, cells were grown overnight aerobically at 30 ºC in 4 ml of LB. The culture vortexed and the OD_600_ measured and set to 100%, then tubes were kept at room temperature standing without agitation for 24h and the OD_600_ of the top part of the culture estimated. To test the ability of biofilm production we used the crystal violet assay ^111^. Briefly, cells were grown aerobically overnight at 30 ºC and 200 µl inoculated into 96-well plates (Thermo Fisher Scientific; MA, USA) at starting OD_600_ of 0.05 and allowed to grow without agitation at room temperature for 6h, 24 h or 4 days. After that, 25 µl of the culture was removed and the OD_600_ measured to estimate the density of planktonic cells. Then, plates were washed with water, stained with 0.1 % (w/v) crystal violet for 30 minutes, and washed again with water. Then, the unwashed dye dissolved with 33 % (v/v) acetic acid and the absorbance at 595 nm measured (biofilm). Biofilm index represents the ratio of biofilm to OD_600_. The permeability of the membrane was visually inspected with the CPRG test described ^81^. Briefly, cells containing the pSEVA2513-LacZ plasmid that constitutively express the β-galactoside (LacZ) enzyme were streaked on an LB agar plates containing 20 µg ml^-1^ of the lactose analogue chlorophenyl red-β-D-galactopyranoside (CPRG), incubated at 30 ºC for 24h or 48h and photographed. To study the influence of NaCl in the growth physiology, we grew both strains in LB without and with different salt concentrations (from 50 mM to 400 mM NaCl) at 30 ºC with 170 rpm. After overnight growth, we directly measured the OD_600_ of cultures (no vortex), then tubes vortexed for 20 seconds to disrupt any flock and measured the OD_600_ (vortex). Then, the ratio OD_600_-vortex / OD_600_-no vortex was calculated and plotted. Bigger numbers for this ratio indicated the presence of bacterial flocks within the liquid media.

### Stress resistance

To evaluate differences in stress resistance between strains we performed filter disc stressor-soaked experiments in solid media. To do that, we added 100 µl of overnight LB grown cultures to pre-warm 0.7 % (w/v) LB top agar, mixed, an overlaid it onto LB agar plates, and let it dry. Then, the filter disc was placed in the middle of the agar plate and soaked with 10 µl of the stressor. After that, the plates were incubated at 30 ºC for 24h, photographed and the diameter of the clear halo generated in both strains compared.

### Protein extraction, western blotting and trypsin digestion

Whole cell protein extracts were prepared collecting cells (OD_600_ ∼1.5) by centrifugation, resuspended in 50 µl of 10 mM Tris-HCl pH 8.0 and 50 µl of 2x SDS-sample buffer (60 µM Tris-HCl pH 6.8, 1 % (w/v) SDS, 5 % (v/v) glycerol, 0.005 % (w/v) bromophenol blue and 1% (v/v) 2-mercaptoethanol) and boiled for 15 minutes. Then, samples were sonicated and cell debris eliminated by centrifugation for 5 minutes at 14.000 x*g*. Supernatants were analyzed on 10 % (w/v) SDS-PAGE gels with a Miniprotean III electrophoresis system (Bio-Rad; CA, USA). For western-blots, proteins were transferred to a polyvinylidene difluoride membrane (Immobilon-P; Merck Millipore, MA, USA) using a Trans-Blot^®^ SD semi-dry transfer cell (Bio-Rad; CA, USA) after denaturing electrophoresis. Membranes were first blocked with PBS buffer containing 3 % (w/v) skimmed milk for 1h at room temperature. Then, membranes were incubated for 1h at room temperature in the same buffer with a 1/2000 dilution of the monoclonal anti-E-tag (Phadia, Sweden) antibody. After that, membranes were washed three times with PBS with 3 % (w/v) skimmed milk and 0.1 % (v/v) Tween-20 to remove the unbound antibody. Next, a 1/5000 dilution of anti-mouse IgG conjugated with peroxidase (POD; Merck, MO, USA) was used to reveal the presence of bound anti-E-tag. The light signal was developed by soaking membranes in BM Chemiluminescence Western blotting substrate (POD; Merck, MO, USA) for 1 minute in the dark and either exposed to an X-ray film or scanned in an Amersham Imager 600 (GE Healthcare, IL, USA). Protease accessibility assays were performed as follows, induced cells were harvested by centrifugation at 4000 x*g* for 3 minutes and resuspended in 100 µl of 10 mM Tris-HCl pH 8.0. Then, this bacterial suspension incubated with 200 µg ml^-1^ trypsin for 20 minutes at 37 ºC. Subsequently, 5 µg ml^-1^ of trypsin inhibitor was added to stop the proteolysis reaction. Finally, samples were centrifugated at 14,000 x*g* for 1 minute and the pellet resuspended in 50 µl of 10 mM Tris-HCl pH 8.0. Then, whole proteins extracted and analyzed by western blot as described earlier. Bacterial permeabilization was performed by adding 400 µl lysozyme (20 mg ml^-1^) to a 1 ml of induced cells and incubated for 2 minutes at RT. Then, cells were washed with 1x PBS and treated with trypsin as described above and the presence of the recombinant protein analyzed by western blot. Also, permeabilized bacteria were visualized with fluorescence microscopy by incubating cells for 30 min in PBS with 3 % (w/v) bovine serum albumin. Then, incubated with the anti-E-tag antibody (1:50) at 4 ºC. After that, samples were washed three times with PBS and incubated with anti-mouse IgG Alexa Fluor 594 (1:500; ThermoFisher Scientific) for 30 minutes in the dark at RT. Finally, cells were visualized in Leica DMI600 B fluorescence microscope.

### Characterization of surface exposed proteins

To identify surface exposed proteins, we followed the protocol described in ^67^. Briefly, first nanoparticles were prepared and activated. In order to bind surface exposed proteins to activated nanoparticles (NPs), *P. putida* cells were grown on LB agar plates overnight and scraped them using a sterile loop, then washed with PBS and incubated for 5 min at 37 °C under stirring in the presence of activated NPs (0.5 mg ml^-1^). After that, unbound reactive groups on the NPs were blocked with 200 mM Tris-HCl pH 7.4. Then, cells were disrupted in a French press and NPs recovered through a permanent magnet for 1 h at 4 °C, and washed, twice with water and twice with 1 M NaCl, to remove non-specifically bound material. To remove further non-covalently bound proteins and fragments of the cell envelope, washed NPs from the binding experiments (NP-Env) were incubated for 1 h at 60 °C in the presence of 1% (w/v) SDS. Proteins that remained covalently bound to the SDS-treated NPs (NP-CbP) were then digested with 40 µg ml^-1^ trypsin to break them into fragments suitable for multidimensional protein identification technology (MudPIT) identification ^67^. The activity of NPs to capture proteins shed spontaneously from intact cells and proteins released by spontaneous cell lysis was evaluated through control experiments (NP-Shed). In these controls, the cells washed with PBS were incubated for 5 min at 37 °C without NPs, removed by tandem centrifugation-filtration, and the resulting supernatants were incubated for 5 min at 37 °C with activated NPs; in this way, NPs could only react with released proteins. NPs were then inactivated, and the proteins covalently bound to NPs were identified by MudPIT analysis. NP-CbP bound by NPs at the cell surface were listed by statistical comparison ^67^ of the Spectral Counts (SpC) determined by the MudPIT analyses on NP-Env experiments with those of NP-Shed controls. Two independent experiments, both for NP-Env and NP-Shed, were conducted for each strain. The experimental mass spectra produced by MudPIT analyses were correlated to *in silico* peptide sequences of a non-redundant P. *putida* KT2440 protein database retrieved from NCBI (http://www.ncbi.nlm.nih.gov/). Data processing of raw spectra was performed as described in ^67^.

### Quantification of cellular aggregates

Overnight grown cultures were diluted to an OD_600_ of 0.1, in 100 ml Erlenmeyer flasks containing 10 ml of LB and grown for 2 hours at 30 °C with shaking. Then, cultures were induced overnight by adding 1mM 3-methylbenzoate. After that, the aggregation experiments were prepared as follows: induced cultures were diluted and mixed in 50 ml flask at 1:1 ratio to yield a final OD_600_ of 1 in M9 medium without C-source in a total of 4 ml volume. Then, cells were incubated at RT with moderate orbital shaking (160 rpm) for 30 minutes. After that, 500 µl samples were taken using head-cut tips to minimize aggregate disruption and combined with 500 µl of melted 2 % (w/v) agar M9 without C-source, mixed and 250 µl transferred to a µ-Dish 35 mm (Ibidi; Germany). Samples were inspected using a confocal multispectral Leica TCS SP8 system with a 100/0.5x oil immersion objective, and images were captured using a 3x amplification factor using a Z-step of 0.5 micron and a refresh frequency of 600 Hz. Virtual 3D reconstruction of aggregates was performed using Imaris software v.10 (BITPLANE; UK) by applying a manual thresholding to the raw image to limit the boundaries of every cell and infers its geometric parameters. Then, the following values were calculated for all detected cells in each fluorescent channel (GFP and mCherry): (i) spatial position (**C**), (ii) axial orientation (**v**_z_) and (iii) bacterial length (L_bact_) and imported as CSV files. The quantification of aggregates was performed in MATLAB (The Mathworks Inc.; USA) by generating a matrix containing the euclidean distance (L^i^_j_) among cells. This matrix was used to stablish potential cell neighbors. Cells were considered as neighbors if L^i^_j_ ≤ L_bact_. All identified potential neighbors were individually inspected with a recurrent depth-first search algorithm using a two spherocylinder contact criterium described in ^112-113^ to confirm bacterial contact. Once confirmed as real neighbors, they were tagged as aggregates. The efficiency of aggregation was estimated as: (i) total raw counting of aggregates and (ii) the total fraction of cells within clusters. The analysis was performed with a minimum of 2 biological repetitions, counting at least 500 individual events (cells or aggregates) in total.

## Associated content

### Supporting information

Supplementary **Table S1**: Genomic coordinates of the 23 deletions introduced in *Pseudomonas putida* KT2440 to construct the naked strain EM371.

Supplementary **Table S2**: Oligonucleotides used in this study.

Supplementary **Table S3**: List of surface-associated proteins identified by activated magnetic nanoparticles.

Supplementary **Table S4**: Bacterial strains and plasmids used in this work.

Supplementary **Figure S1**: Genomic regions deleted in *P. putida* KT2440 to construct the EM371 naked strain.

Supplementary **Figure S2**: Growth profiles of *P. putida* KT2440 and the EM371 naked strain under different metabolic regimes.

Supplementary **Figure S3**: Permeabilization and trypsin treatment of EM371 with the control Trx-G^6^V_HH_ plasmid.

Supplementary **Figure S4:** Expression and localization of the Fos and Jun chimera proteins in *P. putida* KT2440.

Supplementary **Figure S5:** Fluorescent microscopy images of the induced aggregation experiments using EM371 as the bacterial chassis.

Supplementary **Figure S6:** Normalized frequency distribution of cells per aggregate.

## Acknowledgements

Authors are indebted to Maia Kiviisar (Tartu University, Estonia), Raúl Platero (Instituto Clemente Estable, Uruguay), Cristina Patiño (Electron Microscopy Facility CNB, Madrid), Sylvia Gutiérrez (Advanced Light Microscopy, Madrid) and Pavel Dvorak (Masaryk University, Czech Republic) for valuable advice and help with some experiments.

## Author Contributions

EMG, VdL and GB planned the experiments; EMG, SF, DRE and DV did the practical work. All Authors analyzed and discussed the data and contributed to the writing of the article.

## Notes

The authors declare no competing financial interest

## REFERENCES

(1) Hays, S. G.; Patrick, W. G.; Ziesack, M.; Oxman, N.; Silver, P. A., (2015) Better together: engineering and application of microbial symbioses. Curr Opin Biotechnol. 36 (36), 40–49.

(2) McCarty, N. S.; Ledesma Amaro, R., (2019) Synthetic Biology Tools to Engineer Microbial Communities for Biotechnology. Trends Biotechnol. 37 (2), 181–197.

(3) Roell, G. W.; Zha, J.; Carr, R. R.; Koffas, M. A.; Fong, S. S.; Tang, Y. J., (2019) Engineering microbial consortia by division of labor. Microb Cell Fact. 1–11.

(4) Flemming, H.-C.; Wingender, J.; Szewzyk, U.; Steinberg, P.; Rice, S. A.; Kjelleberg, S., (2016) Biofilms: an emergent form of bacterial life. Nat Rev Microbiol. 14 (9), 563–575.

(5) Hall-Stoodley, L.; Costerton, J. W.; Stoodley, P., (2004) Bacterial biofilms: from the Natural environment to infectious diseases. Nat Rev Microbiolo. 2 (2), 95–108.

(6) Davey, M. E.; O’toole, G. A., (2000) Microbial Biofilms: from Ecology to Molecular Genetics. Microbiol Mol Biol Rev. 64, 847–867.

(7) Jenal, U.; Reinders, A.; Lori, C., (2017) Cyclic di-GMP: second messenger extraordinaire. Nat Rev Microbiol. 15 (5), 271–284.

(8) Römling, U.; Galperin, M. Y.; Gomelsky, M., (2013) Cyclic di-GMP: the First 25 Years of a Universal Bacterial Second Messenger. Microbiol Mol Biol Rev. 77, 1–52.

(9) Österberg, S.; Åberg, A.; Herrera Seitz, M. K.; Wolf-Watz, M.; Shingler, V., (2013) Genetic dissection of a motility-associated c-di-GMP signalling protein of *Pseudomonas putida*. Environ Microbiol Rep. 5 (4), 556–565.

(10) Benedetti, I.; de Lorenzo, V.; Nikel, P. I., (2016) Genetic programming of catalytic *Pseudomonas putida* biofilms for boosting biodegradation of haloalkanes. Metab Eng 33, 109–118.

(11) Veiga, E.; de Lorenzo, V.; Fernandez, L. A., (2003) Autotransporters as scaffolds for novel bacterial adhesins: surface properties of *Escherichia coli* cells displaying Jun/Fos dimerization domains. J Bacteriol 185 (18), 5585–5590.

(12) Salema, V.; Marín, E.; Martínez-Arteaga, R.; Ruano-Gallego, D.; Fraile, S.; Margolles, Y.; Teira, X.; Gutiérrez, C.; Bodelón, G.; Fernández, L. A., (2013) Selection of Single Domain Antibodies from Immune Libraries Displayed on the Surface of *E. coli* Cells with Two β-Domains of Opposite Topologies. PLoS ONE 8 (9), e75126.

(13) Piñero-Lambea, C.; Bodelón, G.; Fernández-Periáñez, R.; Cuesta, A. M.; Álvarez-Vallina, L.; Fernández, L. A., (2014) Programming Controlled Adhesion of *E. coli* to Target Surfaces, Cells, and Tumors with Synthetic Adhesins. ACS Synth Biol. 140729085126004.

(14) Glass, D. S.; Riedel-Kruse, I. H., (2018) A Synthetic Bacterial Cell-Cell Adhesion Toolbox for Programming Multicellular Morphologies and Patterns. Cell 174 (3), 649-658.e616.

(15) Nikaido, H.; Vaara, M., (1985) Molecular basis of bacterial outer membrane permeability. Microbiol Rev. 49 (1), 1–32.

(16) Nikaido, H., (2003) Molecular Basis of Bacterial Outer Membrane Permeability Revisited. Microbiol Mol Biol Rev. 67 (4), 593–656.

(17) Tolker-Nielsen, T., (2015) Biofilm Development. Microbiology Spectrum 3 (2), 1–12.

(18) Martinez-Garcia, E.; Nikel, P. I.; Aparicio, T.; de Lorenzo, V., (2014) Pseudomonas 2.0: genetic upgrading of *P. putida* KT2440 as an enhanced host for heterologous gene expression. Microb Cell Fact 13 (1), 159.

(19) Martinez-Garcia, E.; de Lorenzo, V., (2016) The quest for the minimal bacterial genome. Curr Opin Biotechnol 42, 216–224.

(20) de Lorenzo, V., (2008) Systems biology approaches to bioremediation. Curr Opin Biotechnol 19 (6), 579–589.

(21) Kampers, L. F. C.; Volkers, R. J. M.; Martins Dos Santos, V. A. P., (2019) *Pseudomonas putida* KT2440 is HV1 certified, not GRAS. Microb Biotechnol 12 (5), 845–848.

(22) Jimenez, J. I.; Minambres, B.; Garcia, J. L.; Diaz, E., (2002) Genomic analysis of the aromatic catabolic pathways from *Pseudomonas putida* KT2440. Environ Microbiol 4 (12), 824–841.

(23) Kim, J.; Park, W., (2014) Oxidative stress response in *Pseudomonas putida*. App Microbiol Biotechnol 98 (16), 6933–6946.

(24) Nikel, P. I.; Chavarria, M.; Danchin, A.; de Lorenzo, V., (2016) From dirt to industrial applications: *Pseudomonas putida* as a Synthetic Biology chassis for hosting harsh biochemical reactions. Curr Opin Chem Biol 34, 20–29.

(25) Ramos, J.-L.; Sol Cuenca, M.; Molina-Santiago, C.; Segura, A.; Duque, E.; Gómez-García, M. R.; Udaondo, Z.; Roca, A., (2015) Mechanisms of solvent resistance mediated by interplay of cellular factors in *Pseudomonas putida*. FEMS Microbiol Rev 39 (4), 555–566.

(26) Martínez-García, E.; de Lorenzo, V., (2017) Molecular tools and emerging strategies for deep genetic/genomic refactoring of *Pseudomonas*. Curr Opin Biotechnol 47, 120–132.

(27) Martínez-García, E.; Goñi-Moreno, A.; Bartley, B.; McLaughlin, J.; Sánchez-Sampedro, L.; Pascual del Pozo, H.; Prieto Hernández, C.; Marletta, A. S.; de Lucrezia, D.; Sánchez-Fernández, G.; Fraile, S.; de Lorenzo, V., (2019) SEVA 3.0: an update of the Standard European Vector Architecture for enabling portability of genetic constructs among diverse bacterial hosts. Nucleic Acids Res 48(6), 3395.

(28) Nikel, P. I.; Martinez-Garcia, E.; de Lorenzo, V., (2014) Biotechnological domestication of pseudomonads using synthetic biology. Nat Rev Microbiol 12 (5), 368–379.

(29) Poblete-Castro, I.; Becker, J.; Dohnt, K.; dos Santos, V. M.; Wittmann, C., (2012) Industrial biotechnology of *Pseudomonas putida* and related species. Appl Microbiol Biotechnol 93 (6), 2279–2290.

(30) Loeschcke, A.; Thies, S., (2015) *Pseudomonas putida*—a versatile host for the production of natural products. Appl Microbiol Biotechnol 99 (15), 6197–6214.

(31) Nikel, P. I.; de Lorenzo, V., (2018) *Pseudomonas putida* as a functional chassis for industrial biocatalysis: From native biochemistry to trans-metabolism. Metab Eng 50, 142–155.

(32) Valls, M.; de Lorenzo, V.; Gonzàlez-Duarte, R.; Atrian, S., (2000) Engineering outer-membrane proteins in *Pseudomonas putida* for enhanced heavy-metal bioadsorption. J Inorg Biochem 79 (1-4), 219–223.

(33) Winsor, G. L.; Lam, D. K.; Fleming, L.; Lo, R.; Whiteside, M. D.; Yu, N. Y.; Hancock, R. E.; Brinkman, F. S., (2011) *Pseudomonas* Genome Database: improved comparative analysis and population genomics capability for *Pseudomonas* genomes. Nucleic Acids Res. 39 (Database issue), D596–600.

(34) Martínez-García, E.; de Lorenzo, V., (2011) Engineering multiple genomic deletions in Gram-negative bacteria: analysis of the multi-resistant antibiotic profile of *Pseudomonas putida* KT2440. Environ. Microbiol. 13 (10), 2702–2716.

(35) Martínez-García, E.; de Lorenzo, V., (2012) Transposon-based and plasmid-based genetic tools for editing genomes of Gram-negative bacteria. Methods Mol. Biol. 813, 267–283.

(36) Ruer, S.; Stender, S.; Filloux, A.; de Bentzmann, S., (2007) Assembly of fimbrial structures in *Pseudomonas aeruginosa*: functionality and specificity of chaperone-usher machineries. J Bacteriol 189 (9), 3547–3555.

(37) Tomaras, A. P.; Dorsey, C. W.; Edelmann, R. E.; Actis, L. A., (2003) Attachment to and biofilm formation on abiotic surfaces by *Acinetobacter baumannii:* involvement of a novel chaperone-usher pili assembly system. Microbiology 149 (Pt 12), 3473–3484.

(38) Nuccio, S. P.; Baumler, A. J., (2007) Evolution of the chaperone/usher assembly pathway: fimbrial classification goes Greek. Microbiol Mol Biol Rev 71 (4), 551–575.

(39) Burrows, L. L., (2012) *Pseudomonas aeruginosa* twitching motility: type IV pili in action. Annu Rev Microbiol 66, 493–520.

(40) Martínez-García, E.; Nikel, P. I.; Chavarría, M.; de Lorenzo, V., (2014) The metabolic cost of flagellar motion in *Pseudomonas putida* KT2440. Environ. Microbiol. 16 (1), 291–303.

(41) Rocchetta, H. L.; Burrows, L. L.; Lam, J. S., (1999) Genetics of O-antigen biosynthesis in *Pseudomonas aeruginosa*. Microbiol Mol Biol Rev 63 (3), 523–553.

(42) Raetz, C. R. H.; Whitfield, C., (2002) Lipopolysaccharide endotoxins. Annu Rev Biochem 71 (1), 635–700.

(43) Ramos-Gonzalez, M. I.; Ruiz-Cabello, F.; Brettar, I.; Garrido, F.; Ramos, J. L., (1992) Tracking genetically engineered bacteria: monoclonal antibodies against surface determinants of the soil bacterium *Pseudomonas putida* 2440. J Bacteriol 174 (9), 2978–2985.

(44) Junker, F.; Rodríguez-Herva, J.; Duque, E.; Ramos-González, M. x. E. a.; Llamas, M. x. E. a.; Ramos, J., (2001) A WbpL mutant of *Pseudomonas putida* DOT-T1E strain, which lacks the O-antigenic side chain of lipopolysaccharides, is tolerant to organic solvent shocks. Extremophiles 5 (2), 93–99.

(45) Nelson, K. E.; Weinel, C.; Paulsen, I. T.; Dodson, R. J.; Hilbert, H.; Martins dos Santos, V. A. P.; Fouts, D. E.; Gill, S. R.; Pop, M.; Holmes, M.; Brinkac, L.; Beanan, M.; DeBoy, R. T.; Daugherty, S.; Kolonay, J.; Madupu, R.; Nelson, W.; White, O.; Peterson, J.; Khouri, H.; Hance, I.; Chris Lee, P.; Holtzapple, E.; Scanlan, D.; Tran, K.; Moazzez, A.; Utterback, T.; Rizzo, M.; Lee, K.; Kosack, D.; Moestl, D.; Wedler, H.; Lauber, J.; Stjepandic, D.; Hoheisel, J.; Straetz, M.; Heim, S.; Kiewitz, C.; Eisen, J. A.; Timmis, K. N.; Düsterhöft, A.; Tümmler, B.; Fraser, C. M., (2002) Complete genome sequence and comparative analysis of the metabolically versatile *Pseudomonas putida* KT2440. Environ. Microbiol. 4 (12), 799–808.

(46) Espinosa-Urgel, M.; Salido, A.; Ramos, J. L., (2000) Genetic analysis of functions involved in adhesion of *Pseudomonas putida* to seeds. J Bacteriol 182 (9), 2363–2369.

(47) Yousef-Coronado, F.; Travieso, M. L.; Espinosa-Urgel, M., (2008) Different, overlapping mechanisms for colonization of abiotic and plant surfaces by *Pseudomonas putida*. FEMS Microbiol Lett 288 (1), 118–124.

(48) Hinsa, S. M.; Espinosa-Urgel, M.; Ramos, J. L.; O’Toole, G. A., (2003) Transition from reversible to irreversible attachment during biofilm formation by *Pseudomonas fluorescens* WCS365 requires an ABC transporter and a large secreted protein. Mol Microbiol 49 (4), 905–918.

(49) Newell, P. D.; Monds, R. D.; O’Toole, G. A., (2009) LapD is a bis-(3’,5’)-cyclic dimeric GMP-binding protein that regulates surface attachment by *Pseudomonas fluorescens* Pf0-1. Proc Natl Acad Sci USA 106 (9), 3461–3466.

(50) Newell, P. D.; Boyd, C. D.; Sondermann, H.; O’Toole, G. A., (2011) A c-di-GMP effector system controls cell adhesion by inside-out signaling and surface protein cleavage. PLoS Biol 9 (2), e1000587.

(51) Martinez-Gil, M.; Yousef-Coronado, F.; Espinosa-Urgel, M., (2010) LapF, the second largest *Pseudomonas putida* protein, contributes to plant root colonization and determines biofilm architecture. Mol Microbiol 77 (3), 549–561.

(52) Molina, M. A.; Ramos, J. L.; Espinosa-Urgel, M., (2006) A two-partner secretion system is involved in seed and root colonization and iron uptake by *Pseudomonas putida* KT2440. Environ Microbiol 8 (4), 639–647.

(53) Flemming, H.-C.; Neu, T. R.; Wozniak, D. J., (2007) The EPS Matrix: The “House of Biofilm Cells”. J Bacteriol 189, 7945–7947.

(54) Nilsson, M.; Chiang, W. C.; Fazli, M.; Gjermansen, M.; Givskov, M.; Tolker-Nielsen, T., (2011) Influence of putative exopolysaccharide genes on *Pseudomonas putida* KT2440 biofilm stability. Environ Microbiol 13 (5), 1357–1369.

(55) Serra, D. O.; Richter, A. M.; Hengge, R., (2013) Cellulose as an architectural element in spatially structured *Escherichia coli* biofilms. J Bacteriol 195 (24), 5540–5554.

(56) Nielsen, L.; Li, X.; Halverson, L. J., (2011) Cell-cell and cell-surface interactions mediated by cellulose and a novel exopolysaccharide contribute to *Pseudomonas putida* biofilm formation and fitness under water-limiting conditions. Environ Microbiol 13 (5), 1342–1356.

(57) Hay, I. D.; Wang, Y.; Moradali, M. F.; Rehman, Z. U.; Rehm, B. H., (2014) Genetics and regulation of bacterial alginate production. Environ Microbiol 16 (10), 2997–3011.

(58) Chang, W. S.; van de Mortel, M.; Nielsen, L.; Nino de Guzman, G.; Li, X.; Halverson, L. J., (2007) Alginate production by *Pseudomonas putida* creates a hydrated microenvironment and contributes to biofilm architecture and stress tolerance under water-limiting conditions. J Bacteriol 189 (22), 8290–8299.

(59) Svenningsen, N. B.; Martinez-Garcia, E.; Nicolaisen, M. H.; de Lorenzo, V.; Nybroe, O., (2018) The biofilm matrix polysaccharides cellulose and alginate both protect *Pseudomonas putida* mt-2 against reactive oxygen species generated under matric stress and copper exposure. Microbiology 164 (6), 883–888.

(60) Larsen, P.; Nielsen, J. L.; Dueholm, M. S.; Wetzel, R.; Otzen, D.; Nielsen, P. H., (2007) Amyloid adhesins are abundant in natural biofilms. Environ Microbiol 9 (12), 3077–3090.

(61) Staley, T.; Lawrence, E., (1997) Variable specificity of Tn*7:: lacZY* insertion into the chromosome of root-colonizing *Pseudomonas putida* strains. Molecular Ecology 6, 85–87.

(62) Choi, K. H.; Gaynor, J. B.; White, K. G.; Lopez, C.; Bosio, C. M.; Karkhoff-Schweizer, R. R.; Schweizer, H. P., (2005) A Tn*7*-based broad-range bacterial cloning and expression system. Nat Methods 2 (6), 443–448.

(63) Martinez-Garcia, E.; Jatsenko, T.; Kivisaar, M.; de Lorenzo, V., (2015) Freeing *Pseudomonas putida* KT2440 of its proviral load strengthens endurance to environmental stresses. Environ Microbiol 17 (1), 76–90.

(64) Olaya-Abril, A.; Jimenez-Munguia, I.; Gomez-Gascon, L.; Rodriguez-Ortega, M. J., (2014) Surfomics: shaving live organisms for a fast proteomic identification of surface proteins. J Proteomics 97, 164–176.

(65) Giombini, E.; Orsini, M.; Carrabino, D.; Tramontano, A., (2010) An automatic method for identifying surface proteins in bacteria: SLEP. BMC Bioinformatics 11, 39.

(66) Doro, F.; Liberatori, S.; Rodríguez-Ortega, M. J.; Rinaudo, C. D.; Rosini, R.; Mora, M.; Scarselli, M.; Altindis, E.; D’Aurizio, R.; Stella, M.; Margarit, I.; Maione, D.; Telford, J. L.; Norais, N.; Grandi, G., (2009) Surfome Analysis as a Fast Track to Vaccine Discovery: identification of a novel protective antigen for Group B *Streptococcus* hypervirulent strain COH1. Mol Cell Proteomics 8 (7), 1728–1737.

(67) Vecchietti, D.; Di Silvestre, D.; Miriani, M.; Bonomi, F.; Marengo, M.; Bragonzi, A.; Cova, L.; Franceschi, E.; Mauri, P.; Bertoni, G., (2012) Analysis of *Pseudomonas aeruginosa* cell envelope proteome by capture of surface-exposed proteins on activated magnetic nanoparticles. PLoS One 7 (11), e51062.

(68) Rodriguez-Ortega, M. J.; Norais, N.; Bensi, G.; Liberatori, S.; Capo, S.; Mora, M.; Scarselli, M.; Doro, F.; Ferrari, G.; Garaguso, I.; Maggi, T.; Neumann, A.; Covre, A.; Telford, J. L.; Grandi, G., (2006) Characterization and identification of vaccine candidate proteins through analysis of the group A *Streptococcus* surface proteome. Nat Biotechnol 24 (2), 191–197.

(69) Tjalsma, H.; Lambooy, L.; Hermans, P. W.; Swinkels, D. W., (2008) Shedding & shaving: disclosure of proteomic expressions on a bacterial face. Proteomics 8 (7), 1415–1428.

(70) Tjalsma, H.; Scholler-Guinard, M.; Lasonder, E.; Ruers, T. J.; Willems, H. L.; Swinkels, D. W., (2006) Profiling the humoral immune response in colon cancer patients: diagnostic antigens from *Streptococcus bovis*. Int J Cancer 119 (9), 2127–2135.

(71) Tjalsma, H.; Lasonder, E.; Schöller-Guinard, M.; Swinkels, D. W., (2007) Shotgun immunoproteomics to identify disease-associated bacterial antigens: Application to human colon cancer. PROTEOMICS – Clinical Applications 1 (4), 429–434.

(72) Harshey, R. M., (2003) Bacterial motility on a surface: many ways to a common goal. Annu Rev Microbiol 57, 249–273.

(73) Espeso, D. R.; García, E. M., (2016) Physical forces shape group identity of swimming *Pseudomonas putida* cells. Front Microbiol 7 1437.

(74) Romling, U.; Bian, Z.; Hammar, M.; Sierralta, W. D.; Normark, S., (1998) Curli fibers are highly conserved between Salmonella typhimurium and *Escherichia coli* with respect to operon structure and regulation. J Bacteriol 180 (3), 722–731.

(75) Friedman, L.; Kolter, R., (2003) Genes involved in matrix formation in *Pseudomonas aeruginosa* PA14 biofilms. Mol Microbiol 51 (3), 675–690.

(76) Pesavento, C.; Becker, G.; Sommerfeldt, N.; Possling, A.; Tschowri, N.; Mehlis, A.; Hengge, R., (2008) Inverse regulatory coordination of motility and curli-mediated adhesion in *Escherichia coli*. Genes Dev. 22 (17), 2434–2446.

(77) Serra, D. O.; Richter, A. M.; Klauck, G.; Mika, F.; Hengge, R., (2013) Microanatomy at Cellular Resolution and Spatial Order of Physiological Differentiation in a Bacterial Biofilm. mBio 4.

(78) Rosenberg, M.; Gutnick, D.; Rosenberg, E., (1980) Adherence of bacteria to hydrocarbons: A simple method for measuring cell-surface hydrophobicity. FEMS Microbiology Letters 9 (1), 29–33.

(79) Rosenberg, M., (1984) Bacterial adherence to hydrocarbons: a useful technique for studying cell surface hydrophobicity. FEMS Microbiol Letters 22 (3), 289–295.

(80) Lahesaare, A.; Ainelo, H.; Teppo, A.; Kivisaar, M.; Heipieper, H. J.; Teras, R., (2016) LapF and Its Regulation by Fis Affect the Cell Surface Hydrophobicity of *Pseudomonas putida*. PLoS One 11 (11), e0166078.

(81) Paradis-Bleau, C.; Kritikos, G.; Orlova, K.; Typas, A.; Bernhardt, T. G., (2014) A genome-wide screen for bacterial envelope biogenesis mutants identifies a novel factor involved in cell wall precursor metabolism. PLoS Genet 10 (1), e1004056.

(82) Delcour, A. H., (2009) Outer membrane permeability and antibiotic resistance. Biochim Biophys Acta 1794 (5), 808–816.

(83) Rubin, A. J.; Hayden, P. L.; Hanna, G. P., (1969) Coagulation of *Escherichia coli* by neutral salts. Water Research 3 (11), 843–852.

(84) Lindahl, M.; Faris, A.; Wadstrom, T.; Hjerten, S., (1981) A new test based on ‘salting out’ to measure relative surface hydrophobicity of bacterial cells. Biochim Biophys Acta 677 (3-4), 471–476.

(85) Arakawa, T.; Timasheff, S. N., (1984) Mechanism of protein salting in and salting out by divalent cation salts: balance between hydration and salt binding. Biochemistry 23 (25), 5912–5923.

(86) Israelachvili, J.; Wennerstrom, H., (1996) Role of hydration and water structure in biological and colloidal interactions. Nature 379 (6562), 219–225.

(87) Marrink, S. J.; Berkowitz, M.; Berendsen, H. J. C., (1993) Molecular dynamics simulation of a membrane/water interface: the ordering of water and its relation to the hydration force. Langmuir 9 (11), 3122–3131.

(88) Watanabe, N.; Suga, K.; Umakoshi, H., (2019) Functional Hydration Behavior: Interrelation between Hydration and Molecular Properties at Lipid Membrane Interfaces. Journal of Chemistry 2019, 4867327.

(89) O’Toole, G. A.; Kolter, R., (1998) Flagellar and twitching motility are necessary for *Pseudomonas aeruginosa* biofilm development. Mol Microbiol 30 (2), 295–304.

(90) Pratt, L. A.; Kolter, R., (1998) Genetic analysis of *Escherichia coli* biofilm formation: roles of flagella, motility, chemotaxis and type I pili. Mol Microbiol 30 (2), 285–293.

(91) Sauer, K.; Cullen, M. C.; Rickard, A. H.; Zeef, L. A. H.; Davies, D. G.; Gilbert, P., (2004) Characterization of nutrient-induced dispersion in *Pseudomonas aeruginosa* PAO1 biofilm. J Bacteriol 186 (21), 7312–7326.

(92) Tolker-Nielsen, T.; Brinch, U. C.; Ragas, P. C.; Andersen, J. B.; Jacobsen, C. S.; Molin, S., (2000) Development and Dynamics of *Pseudomonas* sp. Biofilms. J Bacteriol 182 (22), 6482–6489.

(93) Abate, C.; Luk, D.; Gentz, R.; Rauscher, F. J.; Curran, T., (1990) Expression and purification of the leucine zipper and DNA-binding domains of Fos and Jun: both Fos and Jun contact DNA directly. Proc Natl Acad Sci USA 87 (3), 1032–1036.

(94) Jimenez, J. I.; Fraile, S.; Zafra, O.; de Lorenzo, V., (2015) Phenotypic knockouts of selected metabolic pathways by targeting enzymes with camel-derived nanobodies (V(HH)s). Metab Eng 30, 40–48.

(95) Klauser, T.; Pohlner, J.; Meyer, T. F., (1990) Extracellular transport of cholera toxin B subunit using *Neisseria* IgA protease beta-domain: conformation-dependent outer membrane translocation. EMBO J 9 (6), 1991–1999.

(96) Besingi, R. N.; Clark, P. L., (2015) Extracellular protease digestion to evaluate membrane protein cell surface localization. Nat Protoc 10 (12), 2074–2080.

(97) Salema, V.; Fernandez, L. A., (2017) *Escherichia coli* surface display for the selection of nanobodies. Microb Biotechnol 10 (6), 1468–1484.

(98) Schuurmann, J.; Quehl, P.; Festel, G.; Jose, J., (2014) Bacterial whole-cell biocatalysts by surface display of enzymes: toward industrial application. Appl Microbiol Biotechnol 98 (19), 8031–8046.

(99) Wernerus, H.; Stahl, S., (2004) Biotechnological applications for surface-engineered bacteria. Biotechnol Appl Biochem 40 (Pt 3), 209–228.

(100) Gilbert, C.; Ellis, T., (2019) Biological Engineered Living Materials: Growing Functional Materials with Genetically Programmable Properties. ACS Synth Biol 8 (1), 1–15.

(101) Heras, B.; Totsika, M.; the, K. P. P. o., (2014) The antigen 43 structure reveals a molecular Velcro-like mechanism of autotransporter-mediated bacterial clumping. Proc Natl Acad Sci USA 111 (1), 457–462.

(102) Dvorak, P.; Bayer, E.; de Lorenzo, V., (2020) Surface display of designer protein scaffolds on genome-reduced strains of *Pseudomonas putida*. bioRxiv, 2020.2005.2013.093500.

(103) Sambrook, J.; Maniatis, T.; Fritsch, E. F., (1989) Molecular cloning a laboratory manual. Cold Spring Harbor Laboratory Press: Cold Spring Harbor, N.Y.

(104) Koch, B.; Jensen, L. E.; Nybroe, O., (2001) A panel of Tn*7-*based vectors for insertion of the *gfp* marker gene or for delivery of cloned DNA into Gram-negative bacteria at a neutral chromosomal site. J. Microbiol. Methods 45 (3), 187–195.

(105) Rochat, L.; Pechy-Tarr, M.; Baehler, E.; Maurhofer, M.; Keel, C., (2010) Combination of fluorescent reporters for simultaneous monitoring of root colonization and antifungal gene expression by a biocontrol *pseudomonad* on cereals with flow cytometry. Mol Plant Microbe Interact 23 (7), 949–961.

(106) Horton, R. M.; Hunt, H. D.; Ho, S. N.; Pullen, J. K.; Pease, L. R., (1989) Engineering hybrid genes without the use of restriction enzymes: gene splicing by overlap extension. Gene 77 (1), 61–68.

(107) Wong, S. M.; Mekalanos, J. J., (2000) Genetic footprinting with mariner-based transposition in *Pseudomonas aeruginosa*. Proc. Natl. Acad. Sci. USA 97 (18), 10191–10196.

(108) Silva-Rocha, R.; Martínez-García, E.; Calles, B.; Chavarría, M.; Arce-Rodríguez, A.; de Las Heras, A.; Páez-Espino, A. D.; Durante-Rodríguez, G.; Kim, J.; Nikel, P. I.; Platero, R.; de Lorenzo, V., (2012) The Standard European Vector Architecture (SEVA): a coherent platform for the analysis and deployment of complex prokaryotic phenotypes. Nucleic Acids Res 41 (Database issue), D666–675.

(109) Martinez-Garcia, E.; Aparicio, T.; Goni-Moreno, A.; Fraile, S.; de Lorenzo, V., (2015) SEVA 2.0: an update of the Standard European Vector Architecture for de-/re-construction of bacterial functionalities. Nucleic Acids Res 43 (Database issue), D1183–1189.

(110) Hitchcock, P. J.; Brown, T. M., (1983) Morphological heterogeneity among *Salmonella* lipopolysaccharide chemotypes in silver-stained polyacrylamide gels. J Bacteriol 154 (1), 269–277.

(111) O’Toole, G. A., (2011) Microtiter dish biofilm formation assay. J Vis Exp: JoVE (47), 2437.

(112) Vega, C.; Lago, S., (1994) A fast algorithm to evaluate the shortest distance between rods. Computers & Chemistry 18 (1), 55–59.

(113) Pournin, L.; Weber, M.; Tsukahara, M.; Ferrez, J.-A.; Ramaioli, M.; Liebling, T. M., (2005) Three-dimensional distinct element simulation of spherocylinder crystallization. Granular Matter 7 (2), 119–126.

