## Supplementary Figures for "The naked cell: emerging properties of a surfome-streamlined *Pseudomonas putida* strain"

**Supplementary Figure S1.** Genomic regions deleted in *P. putida* KT2440 to construct the EM371 naked strain.

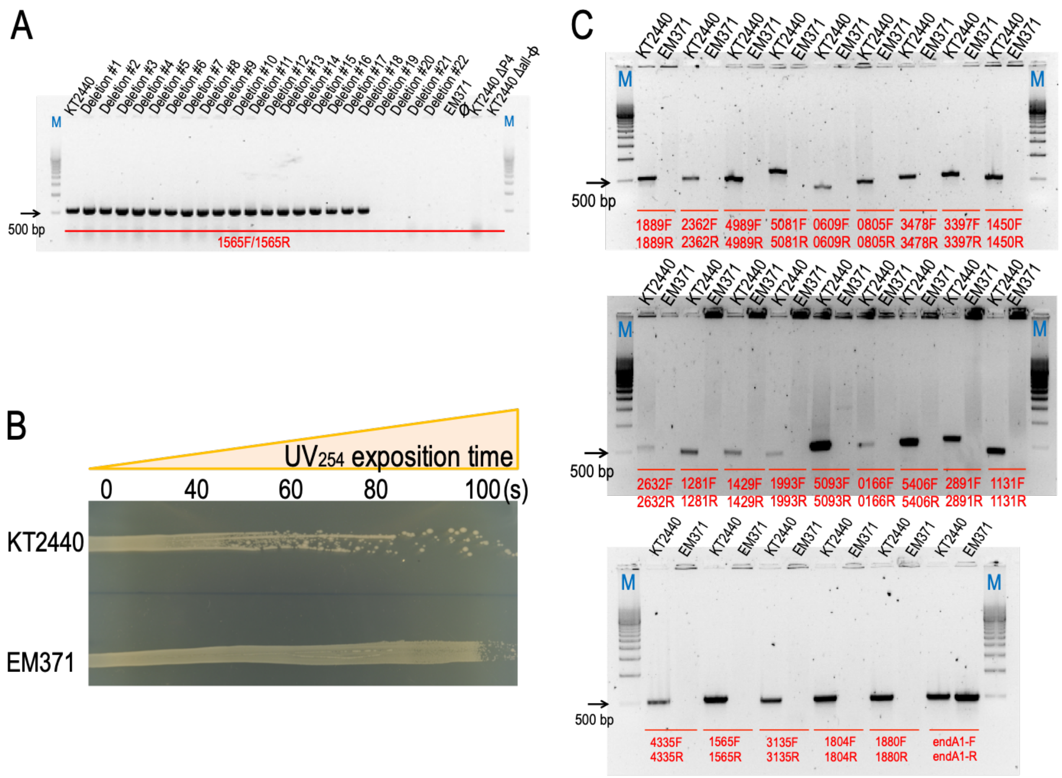


(A) Agarose gel electrophoresis of the diagnostic PCR to identify the deletion step at which the spontaneous excision of prophage 4 occurred {Martinez-Garcia, 2015 #34}. PCR products from different intermediate deletion steps and KT2440 ΔP4 and Δall-ϕ strains as negative controls indicated on top. To confirm the absence of P4 we used the oligos 1565F and 1565R (Table S2) that produced a ~500 bp in strains harboring the prophage. The symbol Ø refers to a blank sample without template. (B) Qualitative UV resistance test. For this experiment, cells were spread onto an LB agar plate and irradiated with UV light for the time indicated. (C) electrophoresis of the diagnostic PCRs to verify the different genomic regions eliminated. Analytical PCR amplifications using either *P. putida* KT2440 or EM371 as DNA template. Oligos, shown in red in the gel, were designed to hybridize within an internal deleted gene (Table S2). Therefore, amplifying only in the wild type a ~500 bp DNA fragment. The oligos endA1-F/endA1-R were used as a *P. putida* control since amplify a fragment of a non-deleted gene in EM371. M corresponds to the 500-bp Molecular Ruler *EZ* load™ (Bio-Rad, Berkeley, CA, USA).

**Supplementary Figure S2.** Growth profiles of *P. putida* KT2440 and the EM371 naked strain under different metabolic regimes.

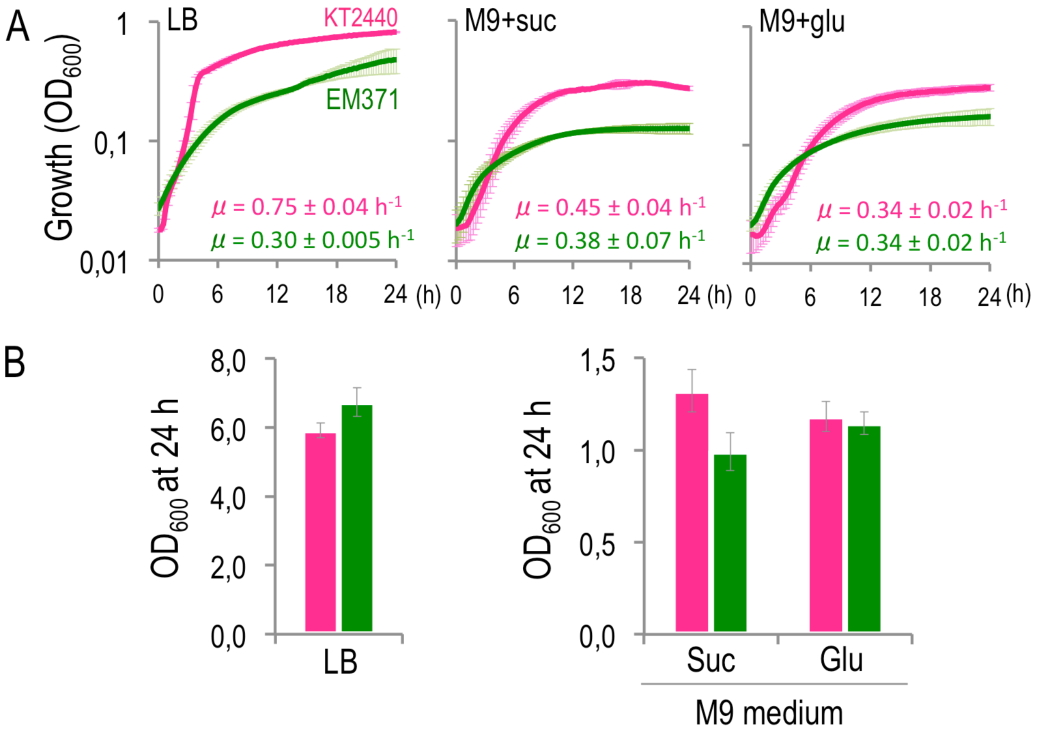


(A) growth curves on rich media (LB), M9 minimal medium with either 0.2 % (w/v) succinate (gluconeogenic condition) or glucose (glycolytic) as sole carbon source. The experiment was performed in Spectramax M² using a 96-well plates in which the $OD_{600}$ was monitored through 24h. The parental strain is depicted in pink while the EM371 in green. The specific growth rates (***μ***) of both strains are located within each plot. (B) end-point cell density ($OD_{600}$) values of the parental (KT2440 in pink) and EM371 (green) strains taken after 24 h of growth in shaken flasks at 30 ºC in LB and M9 with either succinate of glucose as sole carbon source. The average and standard deviation of three independent experiments are shown.

**Supplementary Figure S3.** Permeabilization and trypsin treatment of EM371 with the control Trx-$G^6V_{HH}$ plasmid.

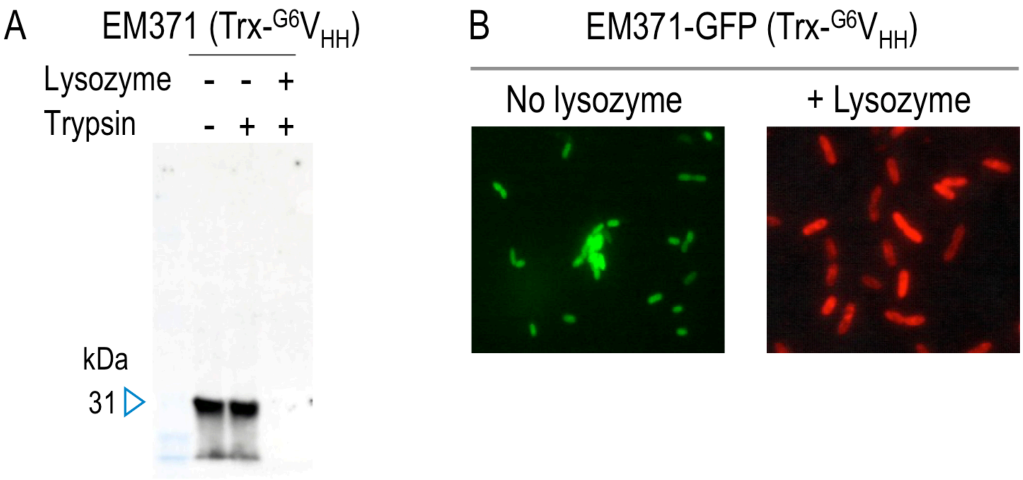


(A) induced cells were permeabilized (+) or not (-) with lysozyme. Following that, samples were incubated (+) or not (-) with trypsin. Then, digestion of the cytoplasmic protein was monitored by western blot using the anti-E-tag antibody and the anti-mouse IgG conjugated with peroxidase. This experiment allows to confirm that the recombinant control protein (Trx-$G^6V_{HH}$) is effectively degraded by trypsin only when cells are permeabilized with lysozyme. (B) Immunofluorescence experiment to visualize the effect of membrane permeabilization. Induced EM371-GFP tagged cells expressing the cytoplasmic control protein (Trx-$G^6V_{HH}$) were treated with a primary anti-E-tag antibody and then with the anti-mouse IgG Alexa fluor 594. Only in permeabilized cells (+ lysozyme) the cytoplasmic control protein is detected (red) while in non-permeabilized bacteria the antibodies do not enter and only the GFP fluorescence is observed.

**Supplementary Figure S4.** Expression and localization of the Fos and Jun chimera proteins in *P. putida* KT2440.

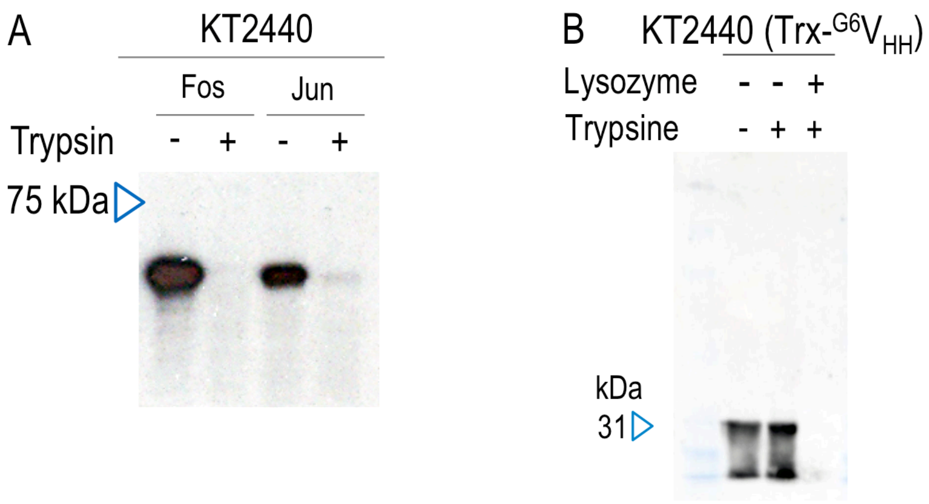


Western blot of induced whole cell extracts containing either pSEVA238-AT-Fos or the pSEVA238-AT-Jun plasmids. Also, induced cells were treated (+) or not (-) with 10 µg ml$^{-1}$ trypsin. The western blot was revealed with anti-E-tag as the primary antibody and reveled with anti-mouse IgG conjugated with peroxidase. (B) permeabilization and trypsin treatment of KT2440 harboring the Trx-G$^{6}$V$_{HH}$ plasmid. Western blot of induced cells permeabilized (+) or not (-) with lysozyme. Then, samples were incubated (+) or not (-) with trypsin.

**Supplementary Figure S5.** Fluorescent microscopy images of the induced aggregation experiments using EM371 as the bacterial chassis.

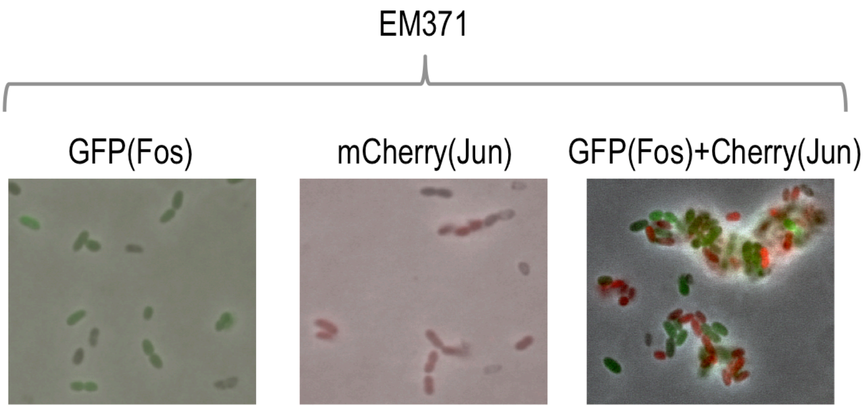


The first image corresponds to EM371-GFP induced cells harboring the hybrid autotransporter with Fos as passenger. The second is induced EM371-mCherry with Jun as passenger. The third one corresponds to a 1:1 aggregation experiment mixing induced cells with the Fos (GFP) and Jun (mCherry) domains and let them stand at room temperature for 20 minutes. Pictures are obtained after merging phase contrast with the fluorescent images of GFP and Red channels.

**Supplementary Figure S6.** Normalized frequency distribution of cells per aggregate.

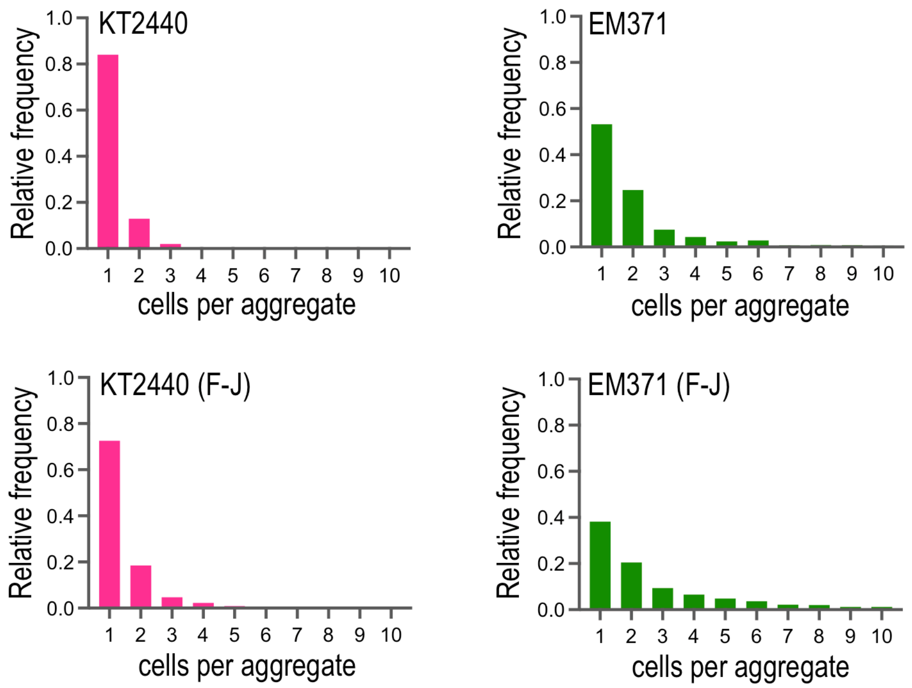


Microscopy images analyzed as described before and the number of cells per aggregate plotted as a normalized histogram. The x-axis was truncated in 10 cells per aggregate for visualization purposes, although the distribution in the case of EM371 (F-J) extends beyond this value (up to 172) in very low frequencies, as can be observed in Fig. 9B.
