## Supplementary Tables S1 S2 S4 for "The naked cell: emerging properties of a surfome-streamlined *Pseudomonas putida* strain"

### SUPPLEMENTARY INFORMATION

**Table S1.** Genomic coordinates of the 23 deletions introduced in *Pseudomonas putida* KT2440 to construct the naked strain EM371.

| Deletion[a] | Gene or gene cluster | Number of genes targeted | Coordinates deleted (bp) | | Extension of the deletion (bp) |
|---|---|---|---|---|---|
| | | | Start | End | |
| 1 | PP1887-PP1891 | 5 | 2,124,627 | 2,133,601 | 8,975 |
| 2 | PP2357-PP2363 | 7 | 2,690,658 | 2,697,068 | 6,411 |
| 3 | PP4986-PP4992 | 7 | 5,680,657 | 5,690,333 | 9,677 |
| 4 | PP5080-PP5083 | 4 | 5,801,586 | 5,805,199 | 3,614 |
| 5 | PP0607-PP0611 | 5 | 715,009 | 717,276 | 2,268 |
| 6 | PP0803-PP0806 | 4 | 921,841 | 945,619 | 23,779 |
| 7 | PP3472-PP3484 | 13 | 3,938,975 | 3,948,636 | 9,662 |
| 8 | PP3396-PP3399 | 4 | 3,846,908 | 3,852,451 | 5,544 |
| 9 | PP1449-PP1450 | 2 | 1,652,238 | 1,658,489 | 6,252 |
| 10 | PP2629-PP2638 | 10 | 3,009,133 | 3,022,978 | 13,846 |
| 11 | PP1277-PP1288 | 12 | 1,458,138 | 1,474,218 | 16,081 |
| 12 | PP1427-PP1430 | 4 | 1,627,269 | 1,631,062 | 3,794 |
| 13 | PP1993 | 1 | 2,259,410 | 2,262,145 | 2,736 |
| 14 | PP5093 | 1 | 5,816,934 | 5,817,944 | 1,012 |
| 15 | PP0164-PP0168 | 5 | 187,239 | 220,542 | 33,304 |
| 16 | PP5404-PP5407 | 4 | 6,161,914 | 6,168,509 | 6,596 |
| 17 | PP2891-PP2893 | 3 | 3,288,245 | 3,294,388 | 6,144 |
| 18 | PP1131 | 1 | 1,294,518 | 1,294,982 | 465 |
| 19 | PP4329-PP4397 | 69 | 4,919,154 | 4,988,328 | 69,175 |
| 20[b] | PP1532-PP1584 | 53 | 1,738,016 | 1,777,398 | 39,383 |
| 21 | PP3132-PP3142 | 11 | 3,547,053 | 3,559,120 | 12,068 |
| 22 | PP1804 | 1 | 2,027,208 | 2,028,212 | 1005 |
| 23 | PP1879-PP1882 | 4 | 2,103,366 | 2,114,282 | 10,917 |

[a] The deletion number identifies the order in which the specific gene or gene cluster was deleted. The information regarding genomic coordinates of each locus was derived from the reported sequence of *P. putida* KT2440 (GenBank #: AE015451) [1].

[b] Spontaneous excision of prophage 4 (PP1532-PP1584), that occurred sometime after deletion #18 [2], during the construction process.

**Table S2.** Oligonucleotides used in this study.

| Name | Sequence (5' → 3')[a] | Usage |
|---|---|---|
| TS1(1887)EcoRI-F | CG**GAATTC**CGGTCTTTTGGCAATTGTCAAT | Deletion of PP1887-PP1891 |
| TS1(1887)-R | TTGGCTCCATCAAAGAATCGAG | Deletion of PP1887-PP1891 |
| TS2(1891)-F | *CTCGATTCTTTGATGGAGCCAA*GACATAACCTG ATTGGGTAAATGGG | Deletion of PP1887-PP1891 |
| TS2(1891)XmaI-R | TCCC**CCCGGG**GGCCAGCGGCTGCACGAGGCC GAAC | Deletion of PP1887-PP1891 |
| TS1(2357)-SacI | ATCC**GAGCTC**CTGCGTAGCTGGGGTAGCCGG | Deletion of PP2357-PP2363 |
| TS1(2357)-R | *TTCGGGGGTATTGACCGCCGT*AAAGCCATTTTT TTTGTATTAGCG | Deletion of PP2357-PP2363 |
| TS2(2363)-F | ACGGCGGTCAATACCCCCGAA | Deletion of PP2357-PP2363 |
| TS2(2363)BamHI-R | CG**GGATCC**CAGCGGCCGCAGTGTCCAG | Deletion of PP2357-PP2363 |
| TS1(4985)-EcoRI-F | CG**GAATTC**ATTCACCGCTGCATGTGCA | Deletion of PP4896-PP4942 |
| TS1(4985)-R | *CAATCCTACTGGTAGGCGTCT*TTAAAAGTCGCC CCACAACTG | Deletion of PP4896-PP4942 |
| TS2(4993)-F | AGACGCCTACCAGTAGGATTG | Deletion of PP4896-PP4942 |
| TS2(4993)BamHI-R | CG**GGATCC**TGCGCATCAGGATCACGTCCAG | Deletion of PP4896-PP4942 |
| TS1(5080)XmaI-F | TCCC**CCCGGG**AGCGCTCGAGAATATCGATCAC C | Deletion of PP5080-PP5083 |
| TS1(5080)-R | *GTAACAGACAGCAAAGGAGTCGCG*TTCTGTGC GAAATTTGATACTTG | Deletion of PP5080-PP5083 |

| | | |
|---|---|---|
| TS2(5083)-F | CGCGACTCCTTTGCTGTCTGTTAC | Deletion of PP5080-PP5083 |
| TS2(5083)BamHI-R | CG**GGATCC**GGGGTCGACGCCGTAGTGGTTGAG | Deletion of PP5080-PP5083 |
| TS1(0607)EcoRI-F | CG**GAATTC**CAGCCGCGTCATCGATGCGCTG | Deletion of PP0607-PP0611 |
| TS1(0607)-R | *TGCACCGGGCATCATTGAACCT*CGCGGTCATTGCAGGAGCGGTC | Deletion of PP0607-PP0611 |
| TS2(0611)-F | AGGTTCAATGATGCCCGGTGCA | Deletion of PP0607-PP0611 |
| TS2(0611)BamHI-R | CG**GGATCC**GGGTTGCGCACATTGGCCACACC | Deletion of PP0607-PP0611 |
| TS1(0803)EcoRI-F | CG**GAATTC**CCAGCACCTGCACCAGGGTGTTG | Deletion of PP0803-PP0806 |
| TS1(0803)-R | *CGGATATCAGCAGGGAGCATCC*TCCAGGCATTCCTGGGTTCTGTG | Deletion of PP0803-PP0806 |
| TS2(0806)-F | GGATGCTCCCTGCTGATATCCG | Deletion of PP0803-PP0806 |
| TS2(0806)XbaI-R | GC**TCTAGA**GATCGGCCGCTCGGCACTCAAGGC | Deletion of PP0803-PP0806 |
| TS1(3472)EcoRI-F | CG**GAATTC**TTCATTTCGGCTAGAAGCAAAGATT | Deletion of PP3472-PP3484 |
| TS1(3472)-R | *GGCAGTATTTGGGAGTCGACCTTG*TTGACTGTTAGGGAAGCTCTGATC | Deletion of PP3472-PP3484 |
| TS2(3484)-F | CAAGGTCGACTCCCAAATACTGCC | Deletion of PP3472-PP3484 |
| TS2(3484)-BamHI-R | CG**GGATCC**CTTCGACAGGGCGCAGTACACTGAA | Deletion of PP3472-PP3484 |
| TS1(3396)EcoRI-F | CG**GAATTC**GTATTCGCTCAGCTCGCTGTCCAG | Deletion of PP3396-PP3399 |

| | | |
|---|---|---|
| TS1(3396)-R | *CACCGCGGTTCAAGGGGGAGCGGT*GAGTCAATTCCTCCTGCAAACGGGC | Deletion of PP3396-PP3399 |
| TS2(3399)-F | ACCGCTCCCCCTTGAACCGCGGTG | Deletion of PP3396-PP3399 |
| TS2(3399)-BamHI-R | CG**GGATCC**GTTGACCAGGCAGGCATCCGGCACG | Deletion of PP3396-PP3399 |
| TS1(1449)EcoRI-F | CG**GAATTC**GGCTACTTCACCGTGCTGACCAATAC | Deletion of PP1449-PP1450 |
| TS1(1449)-R | *CGAGTACGAAAACCGTACATCAT*GACGCCATCGCGGCCCTGGCCATTTG | Deletion of PP1449-PP1450 |
| TS2(1450)-F | ATGATGTACGGTTTTCGTACTCG | Deletion of PP1449-PP1450 |
| TS2(1450)BamHI-R | CG**GGATCC**GCTGAAAACCTACCGCATCTCGCT | Deletion of PP1449-PP1450 |
| TS1(2629)EcoRI-F | CG**GAATTC**TCACCTTGGTAGTGACCCGGCCAGA | Deletion of PP2629-PP2638 |
| TS1(2629)-R | *CCCTTGCAACGCAGCGATTGCCG*TTATACAGCATCAAGGCAGAATGAAATC | Deletion of PP2629-PP2638 |
| TS2(2638)-F | CGGCAATCGCTGCGTTGCAAGGG | Deletion of PP2629-PP2638 |
| TS2(2638)BamHI-R | CG**GGATCC**AAGGCCAGCAACTCACGCAGGTTG | Deletion of PP2629-PP2638 |
| TS1(1277)EcoRI-F | CG**GAATTC**GTCACCGGTGAGTCACTGTGCCAGA | Deletion of PP1277-PP1288 |
| TS1(1277)-R | *CGTGATAAACACATGAGGTGATAGCG*ATGCTGACTCGCCCCTGGGCTGAC | Deletion of PP1277-PP1288 |
| TS2(1288)-F | CGCTATCACCTCATGTGTTTATCACG | Deletion of PP1277-PP1288 |
| TS2(1288)XmaI-R | TCCC**CCCGGG**ATATCTCGCAACAGTTACGTCCTTTA | Deletion of PP1277-PP1288 |

| | | |
|---|---|---|
| TS1(1427)EcoRI-F | CG**GAATTC**TGTTGTCCAGCACTGCAGCGACGC | Deletion of PP1427-PP1430 |
| TS1(1427)-R | *CCCTTTCGCCTTGCGAAACTGC*GAACACTCCTCAGTGAACTCGAAGG | Deletion of PP1427-PP1430 |
| TS2(1430)-F | GCAGTTTCGCAAGGCGAAAGGG | Deletion of PP1427-PP1430 |
| TS2(1430)BamHI-R | CG**GGATCC**TGGGGCAGGTCCATCTTGTTCAGG | Deletion of PP1427-PP1430 |
| TS1(1993)EcoRI-F | CG**GAATTC**GCGCAAGGCCGTGAAGCGGTCAGC | Deletion of PP1993 |
| TS1(1993)-R | *CCAAGAGGCCTGACCTGCTTGCA*GACCTCTTCCCTTGTATGAATCGTC | Deletion of PP1993 |
| TS2(1993)-F | TGCAAGCAGGTCAGGCCTCTTGG | Deletion of PP1993 |
| TS2(1993)BamHI-R | CG**GGATCC**CGAATCGGGTCGTTGTAGATGAC | Deletion of PP1993 |
| TS1(5093)EcoRI-F | CG**GAATTC**AGGCAATGCCGCCAGCAGCCGCGGT | Deletion of PP5093 |
| TS1(5093)-R | *GGCATTCTACCTGCTTGAAGGCCT*TACCTGCAAAGGCCCTTTCGCG | Deletion of PP5093 |
| TS2(5093)-F | AGGCCTTCAAGCAGGTAGAATGCC | Deletion of PP5093 |
| TS2(5093)BamHI-R | CG**GGATCC**CAACTTGCGCAGGCTGGCGAAGGC | Deletion of PP5093 |
| TS1(0164)EcoRI-F | CG**GAATTC**GGATATACCCGCGGGCGCTGGCGAT | Deletion of PP0614-PP0618 |
| TS1(0164)R | *CGCTGTGCGGCCCCGGTGGTCTG*GGCAAAGCCGACATAGCGCACTACGG | Deletion of PP0614-PP0618 |
| TS2(0168)-F | CAGACCACCGGGGCCGCACAGCG | Deletion of PP0614-PP0618 |
| TS2(0168)XmaI-R | TCCC**CCCGGG**CAAGGCGGCTGACATTTTTCACTC | Deletion of PP0614-PP0618 |

| | | |
|---|---|---|
| TS1(2891)EcoRI-F | CG**GAATTC**ACTGGCTTTCCAGCAGTGCCTG | Deletion of PP2891-PP2893 |
| TS1(2891)R | *TTGATGGCCGCCACGCTGCGGGC*TCAGTGCCTTGGAGGTGCCCCAG | Deletion of PP2891-PP2893 |
| TS2(2893)F | GCCCGCAGCGTGGCGGCCATCAA | Deletion of PP2891-PP2893 |
| TS2(2893)BamHI-R | CG**GGATCC**ATTGACCTGCGTGAAATTACC | Deletion of PP2891-PP2893 |
| TS1(1131)EcoRI-F | CG**GAATTC**ACTGCCCTGAGCATTGGGG | Deletion of PP1131 |
| TS1(1131)R | *GGGTTTTTTATTACTGCACTACGG*GGGTAAAGTCTCCATAAGTCAGG | Deletion of PP1131 |
| TS2(1131)F | CCGTAGTGCAGTAATAAAAAACCC | Deletion of PP1131 |
| TS2(1131)BamHI-R | CG**GGATCC**TGCTGGTGCTGGTGGCGATGAA | Deletion of PP1131 |
| TS1(3132)EcoRI-F | CG**GAATTC**GGTACAGCGCCTGGCGCAGCGT | Deletion of PP3132-PP3142 |
| TS1(3132)R | *GCTTGAAGCAGGAGCGCTTCGC*GACTTCAATCTCTGACTGATTGG | Deletion of PP3132-PP3142 |
| TS2(3142)F | GCGAAGCGCTCCTGCTTCAAGC | Deletion of PP3132-PP3142 |
| TS2(3142)BamHI-R | CG**GGATCC**CGAGTTGCGCCTGGTGGCTGAG | Deletion of PP3132-PP3142 |
| TS1(1804)EcoRI-F | CG**GAATTC**CAAGCACGAAGCGGAGCAGGGG | Deletion of PP1804 |
| TS1(1804)R | *GATGCCTTGCGTGCTTGGCCATC*GGGCGGCGTGCACCCAGAAAAG | Deletion of PP1804 |
| TS2(1804)F | GATGGCCAAGCACGCAAGGCATC | Deletion of PP1804 |
| TS2(1804)BamHI-R | CG**GGATCC**GAAATACTGGCGCATGATCAGGCG | Deletion of PP1804 |
| TS1(1879)EcoRI-F | CG**GAATTC**AACCCAGGCTGAATCGTGAGGTT | Deletion of PP1879-PP1882 |

| | | |
|---|---|---|
| TS1(1879)R | *ACAAGCAGAGTGAGAACGTTCATT*CCCGATGCA AAAAAAAAACGCCACC | Deletion of PP1879-PP1882 |
| TS2(1882)F | AATGAACGTTCTCACTCTGCTTGT | Deletion of PP1879-PP1882 |
| TS2(1882)XmaI-R | TCCC**CCCGGG**AACGAAGAAGAAACGACGCTTG GAC | Deletion of PP1879-PP1882 |
| 1889F | GGTAGTTGTCGGCTTCGGTA | Diagnose deletion of PP1887-PP1891 |
| 1889R | TATCGCTATTCGACCCAAGG | Diagnose deletion of PP1887-PP1891 |
| PP2362F | GGACATGCAACTGAGCAAAA | Diagnose deletion of PP2357-PP2363 |
| PP2362R | TCCACACCAGAGAACCACTG | Diagnose deletion of PP2357-PP2363 |
| 4989F | GGCAACATCTTCAGCCTTTC | Diagnose deletion of PP4986-PP4992 |
| 4989R | GATCGACTCGACCATGTCAC | Diagnose deletion of PP4986-PP4992 |
| 5081F | TTAGTGGTGATTGCCGAACA | Diagnose deletion of PP5080-PP5083 |
| 5081R | GCTTGGTGAGGGTGATGAAC | Diagnose deletion of PP5080-PP5083 |
| 0609F | CCAACTTGGTTTTGGTTTGG | Diagnose deletion of PP0607-PP0611 |
| 0609R | ATCGTCCTGGAGCTGGTAGA | Diagnose deletion of PP0607-PP0611 |
| 0805F | GACACAGCCTATGCCTGGAT | Diagnose deletion of PP0803-PP0806 |
| 0805R | CCCCTTGGTAGATGGGAACT | Diagnose deletion of PP0803-PP0806 |

| | | |
|---|---|---|
| 3478F | ATGAAAGAGGGCCAGTACGA | Diagnose deletion of PP3472-PP3484 |
| 3478R | AGATGGTGGCTTTCATGTCC | Diagnose deletion of PP3472-PP3484 |
| 3397F | TCCGATGTGGTACAAGGACA | Diagnose deletion of PP3396-PP3399 |
| 3397R | GTGACCGCTTCGGTGATATT | Diagnose deletion of PP3396-PP3399 |
| 1450F | GTCGAAGGCTTTGTCGAAAC | Diagnose deletion of PP1449-PP1450 |
| 1450R | ACAATACCCGCTCGACACTC | Diagnose deletion of PP1449-PP1450 |
| 2632F | GTTGCTGGTGGGTTACCTGT | Diagnose deletion of PP2629-PP2638 |
| 2632R | GGTAATGTCGCCGATGAAGT | Diagnose deletion of PP2629-PP2638 |
| 1281F | AAGTACGAAGGCTCGGACAA | Diagnose deletion of PP1277-PP1288 |
| 1281R | GCCGACCTTGTATTCCTTCA | Diagnose deletion of PP1277-PP1288 |
| 1429F | GCTGAAACTGATGGGTTGGT | Diagnose deletion of PP1427-PP1430 |
| 1429R | CACTTTGCCCTTGGGTGTAT | Diagnose deletion of PP1427-PP1430 |
| 1993F | TGAAGTCGGCACAGAATCAG | Diagnose deletion of PP1993 |
| 1993R | CCAACCTTCAGCTGGTTGAT | Diagnose deletion of PP1993 |
| 0166F | GATATGGGGCAGTTCAAGGA | Diagnose deletion of PP0164-PP0168 |

| | | |
|---|---|---|
| 0166R | AATTTCGTCCTGCAGTTGCT | Diagnose deletion of PP0164-PP0168 |
| 5406F | CATCTCCTTTCCAACCCAGA | Diagnose deletion of Tn7 |
| 5406R | CGTGCATACCAAACAACAGG | Diagnose deletion of Tn7 |
| 2891F | GCAGGCACTCGGCTACTATC | Diagnose deletion of PP29891-PP2893 |
| 2891R | GTGGTTTACGGGTTTCCAGA | Diagnose deletion of PP29891-PP2893 |
| 1131F | GTAAATCCGCTTTGCTGGTG | Diagnose deletion of PP1131 |
| 1131R | CGGAAGATTTCGTTCTCCTG | Diagnose deletion of PP1131 |
| 4335F | TACCGAGGAACACGAAAACC | Diagnose deletion of the flagellum |
| 4335R | TTGGCAGGTTGTCAGTGAAG | Diagnose deletion of the flagellum |
| 1565F | CTGACCGAGGATCAGATGGT | Diagnose deletion of the prophage 4 |
| 1565R | CCGGGTTGAACTTCACGTAG | Diagnose deletion of the prophage 4 |
| 3135F | CTCAATACCGATGCCTTCGT | Diagnose deletion of PP31321-PP3142 |
| 3135R | TGATGCTTGCGGAAGTACAG | Diagnose deletion of PP31321-PP3142 |
| 1880F | GAGCCCACAATCACCAGTTT | Diagnose deletion of PP1879-PP1882 |
| 1880R | CACCCAGTTCAGTGTCATGG | Diagnose deletion of PP1879-PP1882 |
| 1804F | CGGCAGTGCTGACCAGTGTGTTG | Diagnose deletion of PP1804 |

| | | |
|---|---|---|
| 1804R | CACCCGAAAGTATTAACCACC | Diagnose deletion of PP1804 |
| 5093F | GATCAACCCTCGCTCCCTCAGC | Diagnose deletion of PP5093 |
| 5093R | CCTGGCAACGCTGCTCGACCAG | Diagnose deletion of PP5093 |
| endA1-F | CGCTTTTCGCAGCAGCCTGCCTG | Diagnose presence of *endA-1* |
| endA1-R | GAAGTAGGTGCGGGCGATCATGCC | Diagnose presence of *endA-1* |
| 2357-2363-junction-F | GATACGCTACGCAGCGCAGCAA | Sequence boundaries of PP2353-PP2363 deletion |
| 2357-2363-junction-R | CAGCAGCGCTGGTTCCGTGTG | Sequence boundaries of PP2353-PP2363 deletion |
| Tn7-junction-F | CCGACCTGGGAAGGTCGACTTT | Sequence boundaries of Tn7 deletion |
| Tn7-junction-R | GATGACTTCCTAGGCCATTACTTA | Sequence boundaries of Tn7 deletion |
| Junction Flagella-F | CGCCAAGCCTCGCTACCCGGCCTGCT | Sequence boundaries of flagella deletion |
| Junction Flagella-R | CAGTTGATTCTGGTGGTGCACCCG | Sequence boundaries of flagella deletion |
| curli1-junction-F | CTGCGGTCATCCCAATTAATG | Sequence boundaries of curli operon 1 deletion |
| curli1-junction-R | GGCAGGAAGCGCAACGCCAAG | Sequence boundaries of curli operon 1 deletion |
| curli2-junction-F | GCCACACCACGAACGACATCGG | Sequence boundaries of curli operon 2 deletion |
| curli2-junction-R | CAGACCAGCACATCGCCGTGGC | Sequence boundaries of curli operon 2 deletion |

| | | |
|---|---|---|
| 4986-4992 junction-F | GCACGACCTGCCCGAGGCCCAG | Sequence boundaries of PP4986-PP4992 deletion |
| 4986-4992 junction-R | GGGTCGGGGTGGTGCATTGCG | Sequence boundaries of PP4986-PP4992 deletion |
| 1887-1891 junction-F | ATACCTCGATGGTGCGCTGGGA | Sequence boundaries of PP1887-PP1891 deletion |
| 1887-1891 junction-R | GCAACGGGCCAGTGACCTGCTC | Sequence boundaries of PP1887-PP1891 deletion |
| 2629-2638 junction-F | CAGCACGACCCAGGCAATGAAG | Sequence boundaries of PP2629-PP2638 deletion |
| 2629-2638 junction-R | CGTTCGATGCGTACGCCTGTGC | Sequence boundaries of PP2629-PP2638 deletion |
| 5093-junction-F | CTATTCATAAGAGCTTCATCTATG | Sequence boundaries of PP5093 deletion |
| 5093-junction-R | GCTCATGTCAGGTCCTTGTGGAAA | Sequence boundaries of PP5093 deletion |
| 1993-junction-F | TGCCGGAAGCCAACGCCGAACG | Sequence boundaries of PP1993 deletion |
| 1993-junction-R | TGGCCATGGCGGGTAACACGCAGG | Sequence boundaries of PP1993 deletion |
| 1427-1430 junction-F | GTGAGGTGGAACTCGAAGCCGC | Sequence boundaries of PP1427-PP1430 deletion |
| 1427-1430 junction-R | CAGGTAATTGTCGAACCAAGAGT | Sequence boundaries of PP1427-PP1430 deletion |
| 1277-1288 junction-F | GCCCAGGCCACACACAGCCGCCAG | Sequence boundaries of PP1277-PP1288 deletion |
| 1277-1288 junction-R | CGCGAAGAGGTCAGCCGCCGTGAT | Sequence boundaries of PP1277-PP1288 deletion |
| 5080-5083 junction-F | GATCGGGTAGCTACGCTCGCCCA | Sequence boundaries of PP5080-PP5083 deletion |

| | | |
|---|---|---|
| 5080-5083 junction-R | CAGCTCACGCTCGATTTGCAGGGC | Sequence boundaries of PP5080-PP5083 deletion |
| 1804-junction-F | TGGCCGAGTTCCGTCGGGTGAAT | Sequence boundaries of PP1804 deletion |
| 1804-junction-R | GTCATCATCAATGAACGCCACCG | Sequence boundaries of PP1804 deletion |
| 0607-0611 junction-F | GAAGTCGAAGGCACCATGGGCCA | Sequence boundaries of PP0607-PP0611 deletion |
| 0607-0611 junction-R | TCCAGGCGCCTGCGCTGAGCACA | Sequence boundaries of PP0607-PP0611 deletion |
| 0803-0806 junction-F | GTGTGCACCGACTGCTGCAAGGCC | Sequence boundaries of PP0803-PP0806 deletion |
| 0803-0806 junction-R | TGCACGTGCCACCTTTGCGCGA | Sequence boundaries of PP0803-PP0806 deletion |
| 1449-1450 junction-F | AGCAGCTGTACGTGAAGCTGCAG | Sequence boundaries of PP1449-PP1450 deletion |
| 1449-1450 junction-R | GGCGGCCTTCACCGAAACCATC | Sequence boundaries of PP1449-PP1450 deletion |
| 0164-0168 junction-F | CAGGCCCATGGACGAAAGATGG | Sequence boundaries of PP0164-PP0168 deletion |
| 0164-0168 junction-R | CACATGGTGGTCGAGGTGGGCGC | Sequence boundaries of PP0164-PP0168 deletion |
| 2891-2893 junction-F | GAAGCTGTCTACCAACCCCCAGC | Sequence boundaries of PP2891-PP2893 deletion |
| 2891-2893 junction-R | CGAAATATCCCAGGCGGACAC | Sequence boundaries of PP2891-PP2893 deletion |
| 1131-junction-F | GCCGGGTGCAGAAGGTCACGG | Sequence boundaries of PP1131 deletion |
| 1131-junction-R | CGGCGTCCTGGCCCTGCTCGGC | Sequence boundaries of PP1131 deletion |

| | | |
|---|---|---|
| 3132-3142 junction-F | CGTCGACCAGGGCGCTGTACAA | Sequence boundaries of PP3132-PP3142 deletion |
| 3132-3142 junction-R | CGCCGAGGGGCAACGCCTGGC | Sequence boundaries of PP3132-PP3142 deletion |
| 1879-1882 junction-F | TCACGGCGGTAACAGGGGTTCG | Sequence boundaries of PP1879-PP1882 deletion |
| 1879-1882 junction-R | CCAACTCAGGAAAACCTTGTCA | Sequence boundaries of PP1879-PP1882 deletion |
| 3104-3110 Junction-F | GATGGAAGAGCTGACTTACG | Sequence boundaries of PP3104-PP3110 deletion |
| 3104-3110 Junction-R | CGAGCTCCAGAAAGAAATC | Sequence boundaries of PP3104-PP3110 deletion |
| PP1532-XmaF | TCCC**CCCGGG**GACCAGGCGGTGCGACAGCA | Deletion of prophage 4 |
| PP1586-BamR | CG**GGATCC**CCAACACGAAGCTGAAGCTGGC | Deletion of prophage 4 |
| pEMG-F1 | CCATTCAGGCTGCGCAACTGTTG | To sequence TS1-TS2 in pEMG |
| pEMG-R1 | CTTTACACTTTATGCTTCCGGC | To sequence TS1-TS2 in pEMG |
| pSW-F | GGACGCTTCGCTGAAAACTA | Diagnose curation of the plasmid pSW-I |
| pSW-R | AACGTCGTGACTGGGAAAAC | Diagnose curation of the plasmid pSW-I |

[a] Recognition site for the restriction enzymes specified are indicated in boldface in the DNA sequence, and complementary sequences used in splicing by overlap extension (SOEing) PCR amplifications are shown in italics.

**TABLE S3.** List of surface-associated proteins identified by activated magnetic nanoparticles.

**TABLE S4.** Bacterial strains and plasmids used in this work.

| Strain or plasmid | Relevant characteristics [a] | Reference or source |
|---|---|---|
| *Escherichia coli* | | |
| DH5α λ*pir* | Cloning host; F⁻ λ⁻ *endA1 glnX44*(AS) *thiE1 recA1 relA1 spoT1 gyrA96*($Nal^R$) *rfbC1 deoR nupG* Φ80(*lacZ*Δ*M15*) Δ(*argF-lac*)*U169 hsdR17*($r_K^-$ $m_K^+$) λ*pir* lysogen | 3 |
| HB101 | Helper strain; F⁻ λ⁻ *hsdS20*($r_B^-$ $m_B^-$) *recA13 leuB6*(Am) *araC14* Δ(*gpt-proA*)*62 lacY1 galK2*(Oc) *xyl-5 mtl-1 thiE1 rpsL20*($Sm^R$) *glnX44*(AS) | 4 |
| *Pseudomonas putida* | | |
| KT2440 | Wild-type strain; mt-2 derivative cured of the TOL plasmid pWW0 | 5 |
| EM371 | KT2440 derivative; ΔPP1887-PP1891 ΔPP2357-PP2363 ΔPP4986-PP4992 ΔPP5080-5083 ΔPP0607-0611 ΔPP0803-PP0806 ΔPP3472-3484 ΔPP3396-PP3399 ΔPP1449-PP1450 ΔPP2629-PP2638 ΔPP1277-PP1288 ΔPP1427-PP1430 ΔPP1993 ΔPP5093 ΔPP0164-PP0168 ΔPP5404-PP5407 ΔPP2891-PP2893 ΔPP1131 ΔPP4329-PP4397 Δprophage4 ΔPP3132-PP3142 ΔPP1804 ΔPP1879-PP1882 | This work |
| KT2440-GFP | KT2440 GFP derivative | 6 |
| KT2440-mCherry | KT2440 mCherry derivative | 2 |
| EM371-GFP | EM371 GFP derivative | This work |
| EM371-mCherry | EM371 mCherry derivative | This work |

| Plasmid | Relevant characteristics | Reference or Source |
| --- | --- | --- |
| pRK600 | Helper plasmid used for conjugation; *oriV*(ColE1), RK2(*mob*$^+$ *tra*$^+$); Cm$^R$ | [7] |
| pEMG | Plasmid used for deletions; *oriV*(R6K), *lacZ*$\alpha$ fragment with two flanking I-*Sce*I recognition sites; Km$^R$ | [8] |
| pSW-I | Helper plasmid used for deletions; *oriV*(RK2), *xylS*, P*m*→*I-Sce*I; Ap$^R$ | [9] |
| pEMG-1887 | pEMG bearing a 1.6-kb TS1-TS2 EcoRI-XmaI insert for deleting the PP1887-PP1891 operon | [2] |
| pEMG-R1 | pEMG bearing a 1.6-kb TS1-TS2 Sac*I*-BamHI insert for deleting the PP2357-PP2363 operon | [8] |
| pEMG-4986 | pEMG bearing a 1-kb TS1-TS2 EcoRI-BamHI insert for deleting the PP4986-PP4992 operon | [10] |
| pEMG-5080 | pEMG bearing a 1-kb TS1-TS2 XmaI-BamHI insert for deleting the PP5080-PP5083 operon | This work |
| pEMG-0607 | pEMG bearing a 1-kb TS1-TS2 EcoRI-BamHI insert for deleting the PP0607-PP0611 operon | This work |
| pEMG-0803 | pEMG bearing a 1-kb TS1-TS2 EcoR*I*-XbaI insert for deleting the PP0803-PP0806 operon | This work |
| pEMG-3472 | pEMG bearing a 1-kb TS1-TS2 EcoRI-BamHI insert for deleting the PP3472-PP3484 operon | This work |
| pEMG-298 | pEMG bearing a 1-kb TS1-TS2 EcoRI-BamHI insert for deleting the PP3396-PP3399 operon | [2] |
| pEMG-1449 | pEMG bearing a 1.1-kb TS1-TS2 EcoRI-BamHI insert for deleting the PP1449-PP1450 operon | This work |
| pEMG-286 | pEMG bearing a 1.2-kb TS1-TS2 EcoRI-BamHI insert for deleting the PP2629-PP2638 operon | [2] |
| pEMG-1277 | pEMG bearing a 1.1-kb TS1-TS2 EcoRI-XmaI insert for deleting the PP1277-PP1288 operon | This work |
| pEMG-1427 | pEMG bearing a 1.2-kb TS1-TS2 EcoRI-BamHI insert for deleting the PP1427-PP1430 operon | This work |

| Plasmid | Relevant characteristics | Reference or source |
|---|---|---|
| pEMG-1993 | pEMG bearing a 1.1-kb TS1-TS2 EcoRI-BamHI insert for deleting the PP1993 gene | This work |
| pEMG-5093 | pEMG bearing a 1.1-kb TS1-TS2 EcoRI-BamHI insert for deleting the PP5093 gene | This work |
| pEMG-0164 | pEMG bearing a 1.4-kb TS1-TS2 EcoRI-XmaI insert for deleting the PP0164-PP0168 operon | This work |
| pEMG-Tn7 | pEMG bearing a 1.6-kb TS1-TS2 EcoRI-XmaI insert for deleting the PP5404-PP5407 operon | [11] |
| pEMG-2891 | pEMG bearing a 1.1-kb TS1-TS2 EcoRI-BamHI insert for deleting the PP2891-PP2893 operon | This work |
| pEMG-1131 | pEMG bearing a 1.2-kb TS1-TS2 EcoRI-BamHI insert for deleting the PP1131 gene | This work |
| pEMG-flagella | pEMG bearing a 1.5-kb TS1-TS2 EcoRI-BamHI insert for deleting the PP4329-PP4397 flagellar operon | [12] |
| pEMG-3132 | pEMG bearing a 1.2-kb TS1-TS2 EcoRI-BamHI insert for deleting the PP3132-PP3142 operon | This work |
| pEMG-1804 | pEMG bearing a 1-kb TS1-TS2 EcoRI-BamHI insert for deleting the PP1804 gene | This work |
| pEMG-1879 | pEMG bearing a 1-kb TS1-TS2 EcoRI-XmaI insert for deleting the PP1879-PP1882 operon | This work |
| pSEVA238 | Expression vector; *oriV*(pBBR1); *xylS*-P*m* | [13] |
| pSEVA238-AT-Jun | Expression vector; *oriV*(pBBR1); *xylS*-P*m*→Jun-*igAβ;* $Km^R$ | This work |
| pSEVA238-AT-Fos | Expression vector; *oriV*(pBBR1); *xylS*-P*m*→Fos-*igAβ;* $Km^R$ | This work |

| Plasmid | Relevant characteristics | Reference or source |
|---|---|---|
| pSEVA238-trx-$^{G6}V_{HH}$ | Expression vector; *oriV*(pBBR1); *xylS*-P*m*→ trx-$^{G6}V_{HH}$; $Km^R$ | This work |
| pME9407 | $Ap^R$, $Gm^R$, *oriV*(pUC19), mini-Tn7, $P_{tac}$→*mCherry* | 14 |
| pBK-miniTn*7*-*gfp2* | $Ap^R$, $Cm^R$, $Gm^R$, *oriV*(pUC19), *mob*$^+$, mini-Tn7, $P_{A1/04/03}$→*gfp2* | 15 |
| pUX-BF13 | $Ap^R$; *ori*R6K, *mob*$^+$, provides the Tn*7* transposition function *in trans* | 16 |
| pSEVA2513 | $Km^R$, *oriV*(RSF1010), $P_{EM7}$ | 17 |
| pSEVA2513-lacZ | $Km^R$, *oriV*(RSF1010), $P_{EM7}$→*lacZ* | This work |

[a] Antibiotic markers: Ap, ampicillin; Cm, chloramphenicol; Km, kanamycin; Nal, nalidixic acid; Sm, streptomycin.

[b] Plasmid pEMG-0164 contain a deletion of a thymidine within the TS1-TS2. Since this T is located within an intergenic region between PP0163 and PP0164 (genome coordinate: 187,126) we decided to maintain it. Therefore, the naked strain contains that deletion as well.

#### REFERENCES

(1) Winsor, G. L.; Lam, D. K.; Fleming, L.; Lo, R.; Whiteside, M. D.; Yu, N. Y.; Hancock, R. E.; Brinkman, F. S., (2011) *Pseudomonas* Genome Database: improved comparative analysis and population genomics capability for *Pseudomonas* genomes. *Nucleic Acids Res. 39 (Database issue),* D596-D600.

(2) Martinez-Garcia, E.; Jatsenko, T.; Kivisaar, M.; de Lorenzo, V., (2015) Freeing *Pseudomonas putida* KT2440 of its proviral load strengthens endurance to environmental stresses. *Environ Microbiol 17* (1), 76-90.

(3) Platt, R.; Drescher, C.; Park, S. K.; Phillips, G. J., (2000) Genetic system for reversible integration of DNA constructs and *lacZ* gene fusions into the *Escherichia coli* chromosome. *Plasmid 43 (*1), 12-23.

(4) Boyer, H. W.; Roulland-Dussoix, D., (1969) A complementation analysis of the restriction and modification of DNA in *Escherichia coli. J. Mol. Biol. 41* (3), 459-472.
(5) Bagdasarian, M.; Lurz, R.; Ruckert, B.; Franklin, F. C.; Bagdasarian, M. M.; Frey, J.; Timmis, K. N., (1981) Specific-purpose plasmid cloning vectors. II. Broad host range, high copy number, RSF1010-derived vectors, and a host-vector system for gene cloning in *Pseudomonas*. *Gene 16* (1-3), 237-247.
(6) Espeso, D. R.; García, E. M., (2016) Physical forces shape group identity of swimming Pseudomonas putida cells. *Front Microbiol 7* 1437.
(7) Kessler, B.; de Lorenzo, V.; Timmis, K. N., (1992) A general system to integrate *lacZ* fusions into the chromosomes of Gram-negative eubacteria: regulation of the Pm promoter of the TOL plasmid studied with all controlling elements in monocopy. *Mol. Gen. Genet. 233* (1-2), 293-301.
(8) Martínez-García, E.; de Lorenzo, V., (2011) Engineering multiple genomic deletions in Gram-negative bacteria: analysis of the multi-resistant antibiotic profile of *Pseudomonas putida* KT2440. *Environ. Microbiol. 13* (10), 2702-2716.
(9) Wong, S. M.; Mekalanos, J. J., (2000) Genetic footprinting with mariner-based transposition in *Pseudomonas aeruginosa*. P*roc. Natl. Acad. Sci. USA 97 (*18), 10191-10196.
(10) Martínez-García, E.; de Lorenzo, V., (2012) Transposon-based and plasmid-based genetic tools for editing genomes of Gram-negative bacteria. *Methods Mol. Biol. 813,* 267-283.
(11) Martinez-Garcia, E.; Nikel, P. I.; Aparicio, T.; de Lorenzo, V., (2014) Pseudomonas 2.0: genetic upgrading of *P. putida* KT2440 as an enhanced host for heterologous gene expression. *Microb Cell Fact 13* (1), 159.
(12) Martínez-García, E.; Nikel, P. I.; Chavarría, M.; de Lorenzo, V., (2014) The metabolic cost of flagellar motion in *Pseudomonas putida* KT2440. *Environ. Microbiol. 16* (1), 291-303.
(13) Silva-Rocha, R.; Martínez-García, E.; Calles, B.; Chavarría, M.; Arce-Rodríguez, A.; de Las Heras, A.; Páez-Espino, A. D.; Durante-Rodríguez, G.; Kim, J.; Nikel, P. I.; Platero, R.; de Lorenzo, V., (2012) The Standard European Vector Architecture (SEVA): a coherent platform for the analysis and deployment of complex prokaryotic phenotypes. *Nucleic Acids Res. 41 (Database issue*), D666-675.
(14) Rochat, L.; Pechy-Tarr, M.; Baehler, E.; Maurhofer, M.; Keel, C., (2010) Combination of fluorescent reporters for simultaneous monitoring of root colonization and antifungal gene expression by a biocontrol pseudomonad on cereals with flow cytometry. *Mol Plant Microbe Interact 23* (7), 949-961.

(15) Koch, B.; Jensen, L. E.; Nybroe, O., (2001) A panel of Tn7-based vectors for insertion of the *gfp* marker gene or for delivery of cloned DNA into Gram-negative bacteria at a neutral chromosomal site. J*. Microbiol. Methods 45* (3), 187-195.
(16) Bao, Y.; Lies, D. P.; Fu, H.; Roberts, G. P., (1991) An improved Tn7-based system for the single-copy insertion of cloned genes into chromosomes of gram-negative bacteria. *Gene 109* (1), 167-168.
(17) Martinez-Garcia, E.; Aparicio, T.; Goni-Moreno, A.; Fraile, S.; de Lorenzo, V., (2015) SEVA 2.0: an update of the Standard European Vector Architecture for de-/re-construction of bacterial functionalities. *Nucleic Acids Res 43 (Database issue)*, D1183-1189.
